# Identification of a self-renewing muscle satellite cell state by single-cell chromatin accessibility profiling

**DOI:** 10.1101/2022.09.16.508339

**Authors:** Arinze E. Okafor, Xin Lin, Chenghao Situ, Xiaolin Wei, Xiuqing Wei, Zhenguo Wu, Yarui Diao

## Abstract

A balance between self-renewal and differentiation is critical for the regenerative capacity of tissue-resident stem cells. In skeletal muscle, successful regeneration requires the orchestrated activation, proliferation, and differentiation of muscle satellite cells (MuSCs) that are normally quiescent. A subset of MuSCs undergoes self-renewal to replenish the stem cell pool, but the features that identify and define self-renewing MuSCs remain to be elucidate. Here, through single-cell chromatin accessibility analysis, we reveal the self-renewal versus differentiation trajectories of MuSCs over the course of regeneration in vivo. We identify TGFBR3 as a unique marker of self-renewing MuSCs that can be purified and efficiently contribute to regeneration after transplantation; and we show that SMAD4 and its downstream genes are genetically required for self-renewal *in vivo* by restricting differentiation. Our study unveils the identity and mechanisms of self-renewing MuSCs, while providing a key resource for comprehensive analysis of muscle regeneration.

## INTRODUCTION

In vertebrates, skeletal muscle has robust regenerative ability due to a unique population of muscle-resident stem cells, also called muscle satellite cells (MuSCs) ^1–5^. In mice, MuSCs originate from Pax3/Pax7-positive progenitor cells in the dermomyotome of developing embryos, and they start to occupy their niche between the basal lamina and sarcolemma of myofibers around E16.5 ^6, 7^. After birth, juvenile MuSCs, which remain Pax7+, are proliferative and contribute to postnatal muscle growth. By postnatal day 21, most of these cells become quiescent but can be reactivated upon muscle injury ^8^. During this process, they first exit quiescence to become activated MuSCs (or ASCs), re-enter the cell cycle to proliferate, and generate numerous muscle progenitor cells (MPCs) ^9^. Studies have shown that the activated and proliferative MPCs are heterogeneous in homeostatic and regenerating muscle. That is, whereas some activated and proliferative cells are committed to myogenic differentiation and fuse to form new myofibers, a subpopulation of MPCs are resistant to differentiation cues and have the potential to replenish the MuSCs pool, a process known as self-renewal ^10^. Despite the significance of self-renewal for regeneration, however, the molecular and cellular features that identify and define self-renewing state of MuSCs remain to be elucidate. This critical knowledge gap prevents not only our comprehensive understanding of the biology of muscle regeneration, but also the potential application of deploying the self-renewing state of MuSCs for effective stem cell based therapeutics.

Transplantation experiments have established that the freshly isolated quiescence MuSCs from the uninjured skeletal muscle have a higher capacity to repopulate the stem cell pool *in vivo* than most activated muscle precursor cells (MPCs) or committed myoblasts ^11^. However, due to the intrinsic “quiescence” properties of satellite cells, a key portion of these quiescent cells are not expected to proliferate, making it extremely challenging to expand or manipulate them without impacting their self-renewal properties. As soon as the quiescent MuSC are isolated from their *in vivo* niche, they immediately lose their stem cell potency and are committed to differentiate ^11–18^. The committed muscle precursors can contribute to new myofibers, but are unable to repopulate the in vivo stem cell pool or support long term muscle homeostasis and regeneration ^11^. Therefore, in particular for the purpose of therapeutic strategies, it is important to understand how to identify and isolate these self-renewing cells from regenerating muscle in vivo. Single-cell studies of muscle regeneration at both the transcriptomic and proteomic levels have been reported recently that have indicated details of mechanisms key for MuSC quiescence, stress response, proliferation, and differentiation^19–22^. Yet, the precise molecular features and functions of self-renewing MuSCs, as well as the mechanisms controlling these features and functions, were not fully explored.

Here, we use single-nuclei ATAC-seq (snATAC-seq) to analyze the accessible cis-regulatory elements (cREs) sequences of large numbers of MuSCs cells before and at multiple timepoints after injury during the process of muscle regeneration in adult mice. The cREs, such as enhancers, play a critical role in regulating spatial-temporal gene expression and fine-tuning cellular states ^23^. Using our rich datasets, we have resolved two distinct trajectories for the descendants of quiescent MuSCs, namely myogenic differentiation and self-renewal. By integrating these data with transcriptomic and protein expression analysis, we identify TGFBR3, a co-receptor of TGFβ superfamily, as a specific marker for the self-renewing MuSCs population, allowing us to purify them for in vivo functional transplantation studies. Furthermore, we find that the transcription factor Smad4 and its downstream genes play a critical role in controlling MuSC self-renewal by preventing an alternative differentiation pathway.

## RESULTS

### Temporally resolved single-cell chromatin accessibility atlas of mouse skeletal muscle regeneration

To determine the spatiotemporal changes of gene regulatory mechanisms key for skeletal muscle regeneration, we carried out single-nuclei ATAC-seq (snATAC-seq) analysis using intact nuclei collected from mice lower hindlimb muscles before and after BaCl_2_-induced acute injury at intervals from 6 hours post-injury (hpi) until seven days post-injury (dpi) (**Fig 1A**). We also collected cells from the contralateral uninjured muscle at 6- and 12-hpi to capture the early “alert” states of MuSCs ^24^. All the nuclei were isolated from snap-frozen muscle tissue, thus preserving their in vivo chromatin states without being exposed to stress from tissue dissociation ^15–18^. A total of 114,004 high-quality nuclei were recovered for downstream analysis (**Fig S1A**, Transcription start site (TSS) enrichment > 8, unique fragments per cell > 1000). Using ArchR ^25^, we identified 14 major cell types, namely mature myofibers (**Fig S1B, S1C**), pericytes, fibro-adipogenic progenitor cells (FAPs), endothelial cells, adipocytes, tenocytes, myocytes, MuSCs and MuSCs-derived muscle precursor cells (MuSC/MPC) (**Fig 1B**), and the immune cell populations that can be further classified into macrophages, neutrophils, T cells, B cells, natural killer cells, mast cells, and dendritic cells populations (**Fig S1D, S1E**).

**Figure 1.**
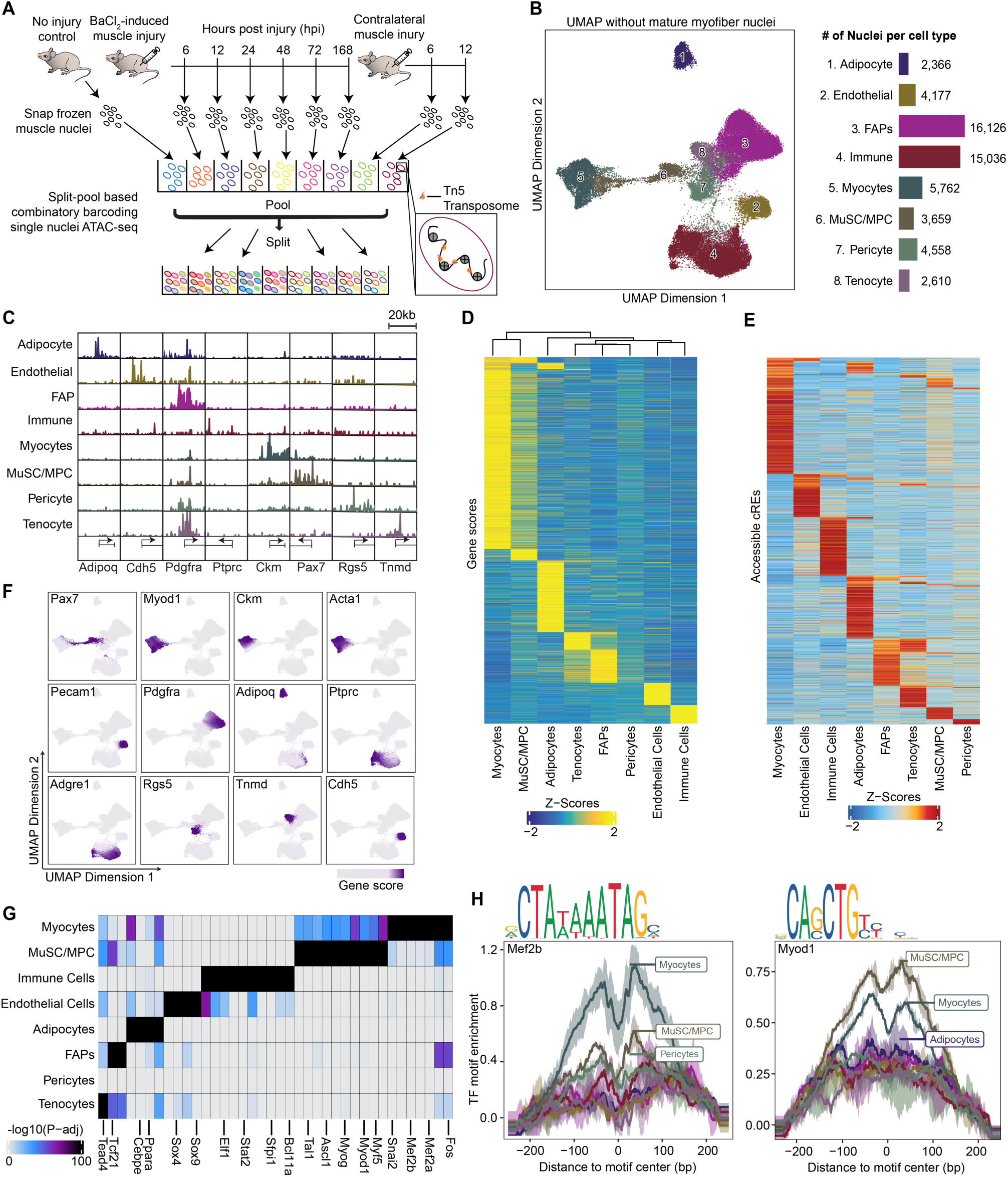
Overview of single-nuclei chromatin accessibility analysis of muscle regeneration. **(A)** Schematic of snATAC-seq experiments. The nuclei were isolated from the snap-frozen intact muscle tissue of lower-hind limb of mice before and after acute injury at the indicated times. Mice for each indicated time: n=2 **(B)** Identification of eight major mononuclear cells types of skeletal muscle before and after injury. After removing the mature myofiber nuclei (**Fig S1C, S1B**), we recovered 54,294 high-quality nuclei (Transcription start site enrichment >8, unique fragments per nuclei > 1,000) from snATAC-seq analysis. **(C)** Genome browser view of chromatin accessibility at selected marker gene loci for each cell type defined in (**B**). **(D)** Heatmap of cell-type specific, highly accessible gene loci quantified by gene scores (FDR <= 0.01, Log2 fold change >= 0.5). Color: scaled gene scores. **(E)** Heatmap of cell-type specific accessible cis-regulatory elements (cREs) (FDR <= 0.01, Log2FC >= 0.5). Color: Z scores of cRE accessibility. **(F)** UMAP showing the gene scores patterns of known marker genes for each cell type. **(G)** In the cell-type specific accessible cREs, transcription factor (TF) motif enrichment was calculated. The significantly enriched TF motif were plotted (FDR <= 0.1, Log2FC >= 0.5) **(H)** TF footprint of Mef2b (left) and Myod1 (right) centered around the accessible cREs in distinct cell types. Top: motif logo.

Because errors in cell assignment may arise in profiling experiments, we assessed chromatin accessibility patterns at specific marker genes known to define each cell type, finding indicators of open chromatin as expected (**Fig 1C**). We also quantified the accessibility of each gene locus as “gene score” ^25^, finding that the gene scores of each cell type strongly correlated with mRNA levels of the matched cell types obtained from published scRNA-seq data of muscle regeneration ^20^ (**Fig S1F)**. Finally, we identified all cell-type-specific genes (based on gene scores), accessible cis-regulatory elements (cRE), and the TF motifs that are significantly enriched on the cREs of specific cell types (**Fig 1D – 1G, Fig S1G, Table S1 – S3**). In total, we cataloged 100,298 accessible cREs across 14 cell types before and after injury at different times of regeneration (**Table S2**). Our data were concordant; for example, the motifs of MEF2B and MYOD1 are most significantly enriched in myocytes and MuSC/MPCs, respectively (**Fig 1H**). Thus, our datasets enable a temporally resolved atlas of chromatin accessibility of MuSCs and their niche populations during regeneration.

### Identification of a self-renewing MuSC population from snATAC-seq analysis of MuSC/MPC

To reveal dynamic changes of MuSC and MuSC-derived MPCs after injury, we next performed a sub-clustering analysis of MuSC/MPCs on a new UMAP space with increased resolution (**Fig 2A)**. We were able to identify five sub-clusters of MuSC/MPCs (**Fig 2A**), including: (1) Clusters 1 and 2 exhibiting high gene scores for *Pax7*, *Cd34*, *Spry1*, *Myf5*, *Foxo1*, *Foxo3, Notch2, Notch3, and Hey1* ^26–30^ (**Fig 2B**); (2) Clusters 4 and 5 showing high gene scores of differentiation makers such as *MyoG*, *Mymk*, and *Ckm* (**Fig 2B**); and (3) Clusters 1 and 3 that are highly proliferative (**Fig 2C**), based on their high chromatin accessibility of a gene list representing features of the G2M phase of the cell cycle ^31^. Therefore, we define the Pax7-high, non-cycling Cluster 2 cells as quiescent MuSCs (QSC); clusters 4 and 5 as two populations of differentiating myoblasts; and cluster 3 as activated/proliferating MPCs that are committed to differentiation due to their low accessibility of *Pax7, Spry1, Notch3*, etc (**Fig 2A, Fig 2B**).

**Figure 2.**
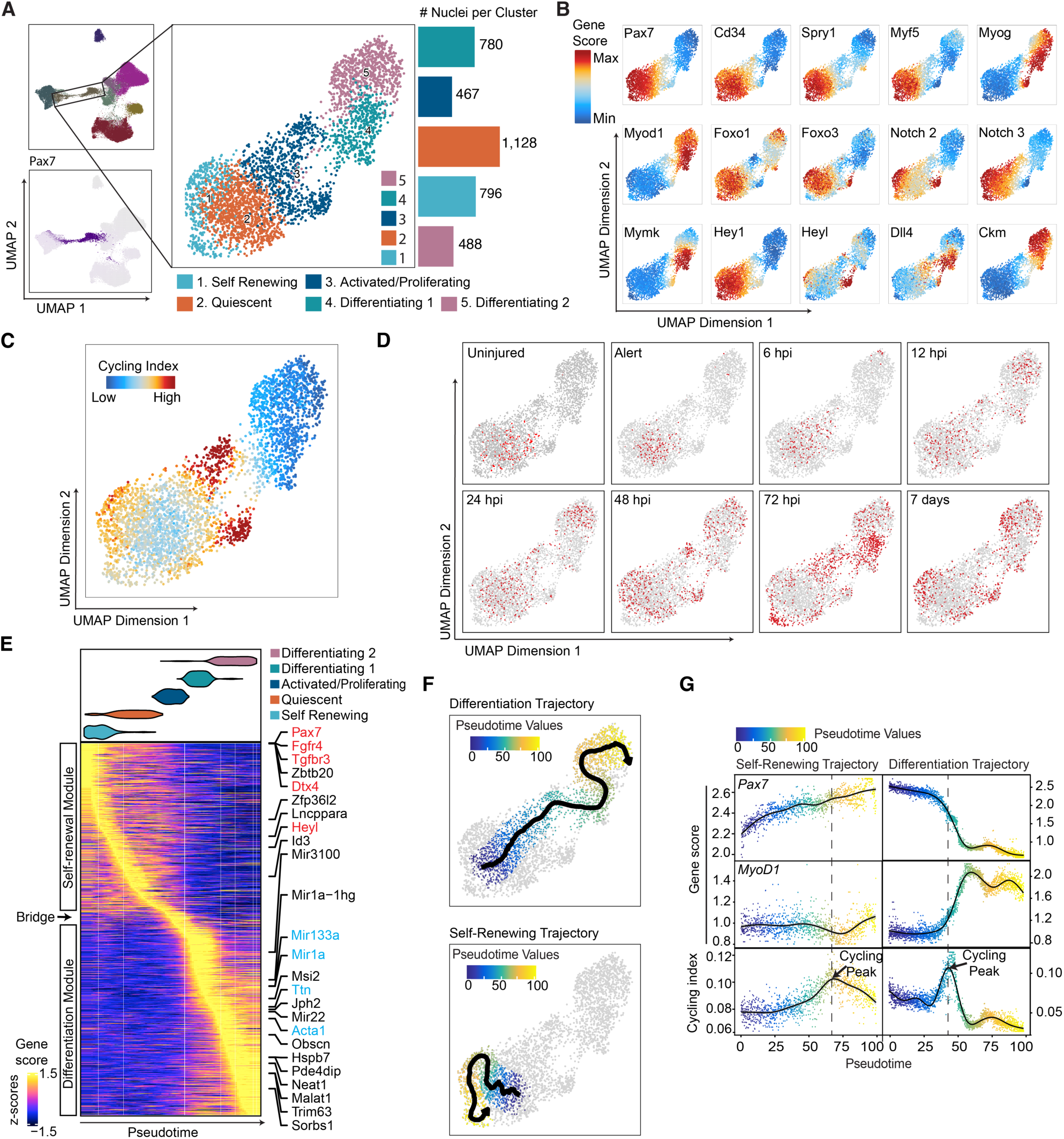
snATAC-seq analysis of MuSC/MPC reveals self-renewal and myogenic differentiation trajectories. **(A)** Five sub-clusters of MuSCs and MuSC-derived muscle progenitor cells (MPC) were plotted on UMAP. **(B)** UMAP plots exhibiting the indicated gene scores across the sub-clusters of MuSC/MPC. **(C)** UMAP plot of cell cycling index across the sub-clusters of MuSC/MPC. The high cell cycling index indicates high proliferative potency. **(D)** The MuSC/MPCs collected at indicated times were plotted as red dots on the UMAP Abbreviations: hpi: hours post-injury; dpi: days post-injury. **(E)** All the MuSC/MPC nuclei are ordered in a pseudotime continuum (x-axis) based on their Euclidean distances in a reduced dimension space. The heatmap shows top-ranked genes co-varying with the pseudotime trajectory (genes as rows, and cells as columns). At the top of the heatmap is a violin plot showing the distribution of cells along the pseudotime for each sub-clusters of MuSC/MPC. Color key: gene scores. **(F)** We set the quiescent MuSC as the cell of origin and starting point of trajectory, and carried out supervised pseudotime trajectory analysis to reveal the myogenic differentiation trajectory (top) and the self-renewal trajectory (bottom). **(G)** Dynamic changes of Pax7 (top) and Myod1 (middle) gene scores, and cell cycling index (bottom) along the self-renewal (left) and the differentiation (right) trajectories. Dashed line indicates the peak of the cycling index for each trajectory.

What is the cellular state of the cells in Cluster 1? The Cluster 1 cells, together with those activated/proliferating cells in Cluster 3, are only present in the injured samples (**Fig 2D**), highly proliferative (**Fig 2C**), but exhibit distinct chromatin accessibility patterns (**Fig 2B**), indicating heterogeneity of MuSC/MPC in the injured muscle. We postulate that the Cluster 1 cells are the candidate self-renewing MuSCs (SRSC), because they are proliferative, high for Pax7/Spry1 gene scores but low for that of MyoD/MyoG. This idea is further supported by the fact that these cycling cells in Cluster 1 are present in late injury timepoint (**Fig 2D**, 48hpi, 72hpi, seven days), but not in samples of uninjured, “Alert”, and early injury timepoints (**Fig 2D,** “uninjured”, “alert”, 6hpi, 12hpi, 24hpi). Taking all these observations into consideration, the cells in Cluster 1 are different from those quiescent ones, or those committed to myogenic differentiation, and likely to be the candidate self-renewing MuSCs.

To delineate the lineage relationship of MuSC/MPCs during regeneration, we carried out trajectory analysis using ArchR ^25^, in which nuclei are ordered in a pseudotime continuum based on their Euclidean distances in a reduced dimension space (**Fig 2E**). Along the pseudotime, we found that quiescence-related genes are most accessible in the early pseudotimes (**Fig 2E**, gene names in red). By contrast, genes and microRNAs ^32^ implicated in myogenic differentiation are more accessible at late pseudotimes (**Fig 2E**, gene names in blue text). Cells ranked along the pseudotime trajectory are clearly separable into two modules based on gene scores, that we name “self-renewal” and “differentiation” (**Fig 2E**). The self-renewal module mainly consists of quiescent and self-renewal clusters, while cells in the differentiation module are well-represented by differentiating myoblasts in Clusters 4 and 5 (**Fig 2E**, top). Notably, the activated/proliferating MuSCs population is ranked in the intermediate “bridge” position connecting the two modules (**Fig 2E**), suggesting that these cells are transitioning from the high stemness, undifferentiated state toward myogenic differentiation.

Given the fact that all the MuSC/MPCs along the pseudotime were derived from QSC, to align this trajectory with stages of regeneration, we designated QSCs as the starting population. This approach identified a myogenic differentiation trajectory and a self-renewing trajectory, both starting from the quiescent MuSCs population (**Fig 2F**). Interestingly, the *Pax7* gene score progressively increases along the self-renewing trajectory but rapidly decreases during differentiation (**Fig 2G,** top). Conversely, while *Myod1* gene score remains low through self-renewal, it increases sharply during myogenic differentiation (**Fig 2G,** middle). We also examined changes in cell cycling indices along the two trajectories (**Fig 2G**, bottom). Our data indicate that both self-renewing and differentiating cells enter cell cycle upon injury, and undergo a transient amplification phase (**Fig 2G**, bottom). These results support the notion that our snATAC-seq analysis of MuSC/MPCs reveals distinct self-renewal and myogenic differentiation trajectories during skeletal muscle regeneration.

### Identification of self-renewing MuSCs population from single-cell RNA-seq analysis

To determine if similar conclusions would emerge from assessment of transcriptome data, we carried out independent single-cell RNA-seq (scRNA-seq) analysis using samples of regenerating skeletal muscle. We selected a timepoint of 3 dpi, as our snATAC-seq data indicated the presence of abundant self-renewing MuSCs at this time (**Fig 2D**, 72dpi). To enrich the MuSC-derived cells away from endothelial and immune cells that are dominant in the injured muscle, we partially depleted CD31+ and CD45+ cells using FACS before scRNA-seq analysis of 3dpi sample ^33^. The cells prepared from the uninjured muscle were also included in the analysis. We identified 15 major cell types from our scRNA-seq data (**Fig S2A, S2B)**, including a population of MuSC/MPCs that can be further classified into four sub-clusters on a new UMAP space (**Fig 3A**). The four sub-populations show distinct expression patterns of *Pax7*, *MyoD1*, proliferation marker *Mki67* (cluster 3) and differentiation marker *Myog* (cluster 4) (**Fig 3B**). We reasoned that the 3pdi, Pax7-high, MyoD-low, and cycling cells in cluster 3 cells (**Fig 3B**, **3C**, the green population) are the candidate self-renewing population. Following the same rationale of our snATAC-seq analysis, we defined the *Myog*-expressing cluster 4 cells as differentiating myoblasts, and the 3dpi cells in cluster 1 as activated MPCs committed to differentiation (activated/committed, **Fig 3C**, bottom, dark red) because they are *Myod1*-high but *Pax7-*low (**Fig 3D**). Taken together, using independent scRNA-seq data, we are able to identify the candidate self-renewing v.s. committed MuSC/MPCs in the injured muscle at 3dpi.

**Figure 3.**
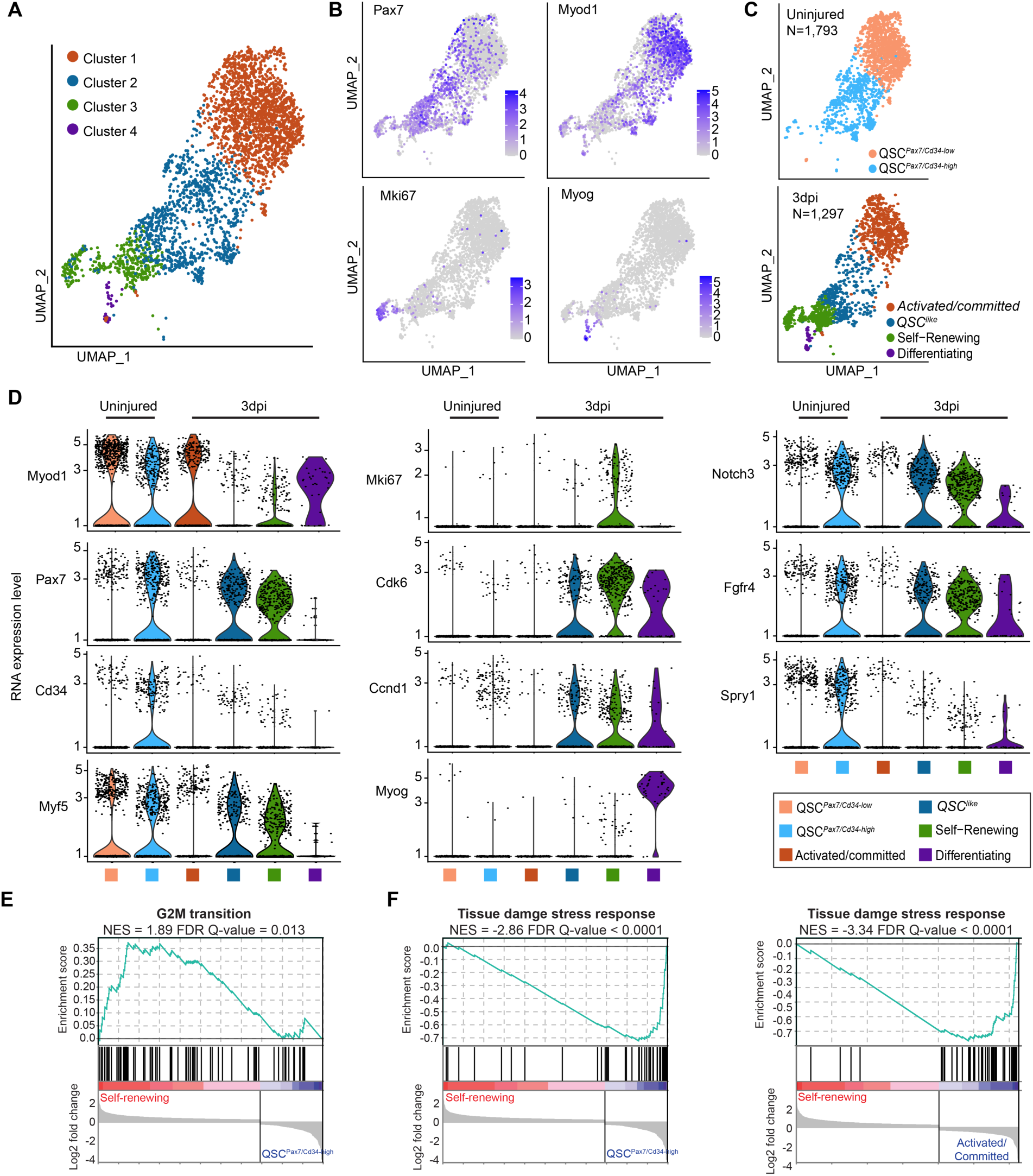
Self-renewing MuSCs exhibit a distinct gene expression pattern compared to the quiescent or committed MuSC/MPCs. **(A)** The MuSC/MPC identified from scRNA-seq analysis of uninjured and 3dpi injured muscle (**Fig S2A, S2B**) were plotted on the UMAP. Four major sub-clusters of MuSC/MPC were identified and plotted in different colors. **(B)** The expression levels of *Pax7*, *Myod1*, *Myog* and *Mki67* mRNAs (log10-normalized) were plotted on the UMAP. Color key: dark color indicates high expression level. **(C)** The MuSC/MPCs shown in **(A)** were separated as uninjured (top) and 3 dpi (bottom), and plotted separately on the same UMPA space shown in (**A**). For each time, the cells were annotated in different colors based on the four major clusters defined in **(A)**. The number of MuSC/MPC for each group: QSC^Pax7/Cd34-low^: 1,181; QSC^Pax7/Cd34-high^: 612; Activated/committed: 533; QSC^like^: 362; Self-Renewing: 366; Differentiating: 36. **(D)** Violin plots showing the mRNA expression levels of the indicated genes across the six sub-clusters defined in (**C**). **(E, F)** Gene set enrichment analysis plots comparing self-renewing MuSCs versus QSC*^Pax7^*^/*Cd34*-high^ or Activated/Committed MPCs. The log2FoldChange for each pairwise differential analysis was used as the rank for GSEA analysis. False Discover Rate (FDR) Q value and Normalized Enrichment Scores (NES) are written on top of each plot. The core stress genes used **(F)** were obtained from Machado et al., 2021^18^.

Notably, the 1,793 uninjured quiescent MuSCs are separated into two sub-clusters from our scRNA-seq analysis (**Fig 3C**, top). A recent study reported that the quiescent MuSC population is composed of “primed” versus “genuine quiescent” MuSCs ^34^, with the latter expressing higher levels of CD34 and Pax7. In support of this observation, we found significantly higher levels of *Cd34* expression in one of the uninjured sub-clusters expressing higher levels of *Pax7* (**Fig 3C, 3D**, pink). Therefore, we classified the quiescent MuSCs into two sub-populations, QSC*^Pax^*^7/*Cd34*- high^ (**Fig 3C, 3D**, pink) versus QSC*^Pax7^*^/*Cd34*-low^ (**Fig 3C**, **3D**, light blue). Recent studies also indicated that *Myod1* transcripts are present at high levels in quiescent MuSCs, despite no detectable MYOD1 protein expression ^35, 36^. Consistent with these studies, all the uninjured quiescent MuSCs express high-levels of *Myod1* RNA (**Fig 3D**, Myod1).

Compared to QSC*^Pax7^*^/*Cd34*-high^, the self-renewing MuSCs (SRSC, **Fig 3D**, green) express low levels of *Cd34*, *Myod1*, and *Spry1*; and high levels of cell cycle genes, including *Mki67*, *Cdk6*, and *Ccnd1* (**Fig 3D**). These differences argue against the possibility that the Pax7+ SRSC are residual QSC remaining from incomplete muscle injury. We also identify 3dpi cells in cluster 2 that we refer to as “QSC-like” MuSCs (QSC^like^) (**Fig 3C**, dark blue). We postulate that these cells are likely self-renewing cells as well, because they express high levels of cell cycling genes (**Fig 3D,** *Cdk6* and *Ccnd1*, despite not expressing *Mki67*) as well as genes preventing the myogenic differentiation fate (**Fig 3D,** *Pax7*, *Notch3*, *Fgfr4*), but low levels of genes that are expressed in QSC such as *Myod1*, *Cd34*, and *Spry1* **(Fig 3D)**. We speculate that the observed differences between the QSC^like^ and SRSC clusters are due at least in part to the fact that the two populations of cells are at different stages of the dynamic self-renewal process.

To further reveal global differences of transcriptome among the sub-populations of MuSC/MPCs, we conducted differential gene expression analysis (**Table S4**) and Gene Set Enrichment Analysis (GSEA) (**Table S5, Fig S2C, S2D**). We found that the genes related to the G2M cell cycle phase are significantly upregulated in both self-renewing (SRSC) and activated/committed MPCs compared to QSC*^Pax7^*^/*Cd34*-high^ (**Fig 3E, Fig S2E**), suggesting their proliferative states at 3dpi upon injury. Furthermore, GSEA results suggest other biological processes, such as extracellular matrix (ECM) degradation and organization, RNA translation, and MAPK signaling, can also distinguish SRSCs from both QSC*^Pax7^*^/*Cd34*-high^ and activated/committed MPCs (**Fig S2C-S2G**). Notably, a recent study had shown that tissue damage induces a conserved cellular stress response that initiates quiescent muscle stem cell activation ^18^. Intriguingly, the expression of these “tissue damage stress response gene sets” ^18^ remains significantly lower in the 3dpi SRSC populations compared to the QSC*^Pax7^*^/*Cd34*-high^ and 3dpi activated/committed MPCs (**Fig 3F**). Together, these data indicate that the SRSC identified from our scRNA-seq analysis is a unique cycling cell state exhibiting distinct gene expression patterns compared to the “quiescence” and the previously defined “activation” state committed to myogenic differentiation.

In total, our observations from scRNA-seq analysis implicate a population of proliferating MuSC/MPCs that are *Pax7*-high, *Myod1*-low, and likely to represent self-renewing MuSCs during regeneration. Importantly, besides having high levels of Pax7 mRNA and proliferative tendencies, these candidate self-renewing MuSCs also exhibit a distinct global gene expression pattern compared to the quiescent MuSCs and the activated/proliferative MPCs committed to differentiation.

### TGFBR3 protein expression identifies self-renewing MuSCs that contribute to the *in vivo* MuSC pool in regeneration

An important goal in the field of muscle regeneration is to identify and isolate the most effective and self-renewable MuSC populations that can support long-term muscle homeostasis and regeneration ^11,37,38^. To identify cell surface markers that enable isolation of self-renewing MuSCs, we reasoned that their expression would correlate with that of *Pax7*, the best-defined MuSC “stemness” marker. We calculated the Pearson’s correlation coefficient (PCC) of mRNA levels and gene scores of all the genes vs *Pax7* across all the MuSC/MPCs. Notably, we found that *Tgfbr3,* which encodes a co-receptor of the TGFβ signaling superfamily and is expressed on the cell surface ^39^, showed a very strong correlation with nascent *Pax7* mRNA and *Pax7* chromatin accessibility levels (**Fig 4A, 4B**). Indeed, we found that *Tgfbr3* gene is highly expressed and highly accessible in the self-renewing MuSC/MPC populations identified from both scRNA-seq and snATAC-seq analysis (**Fig 4C, 4D**).

**Figure 4.**
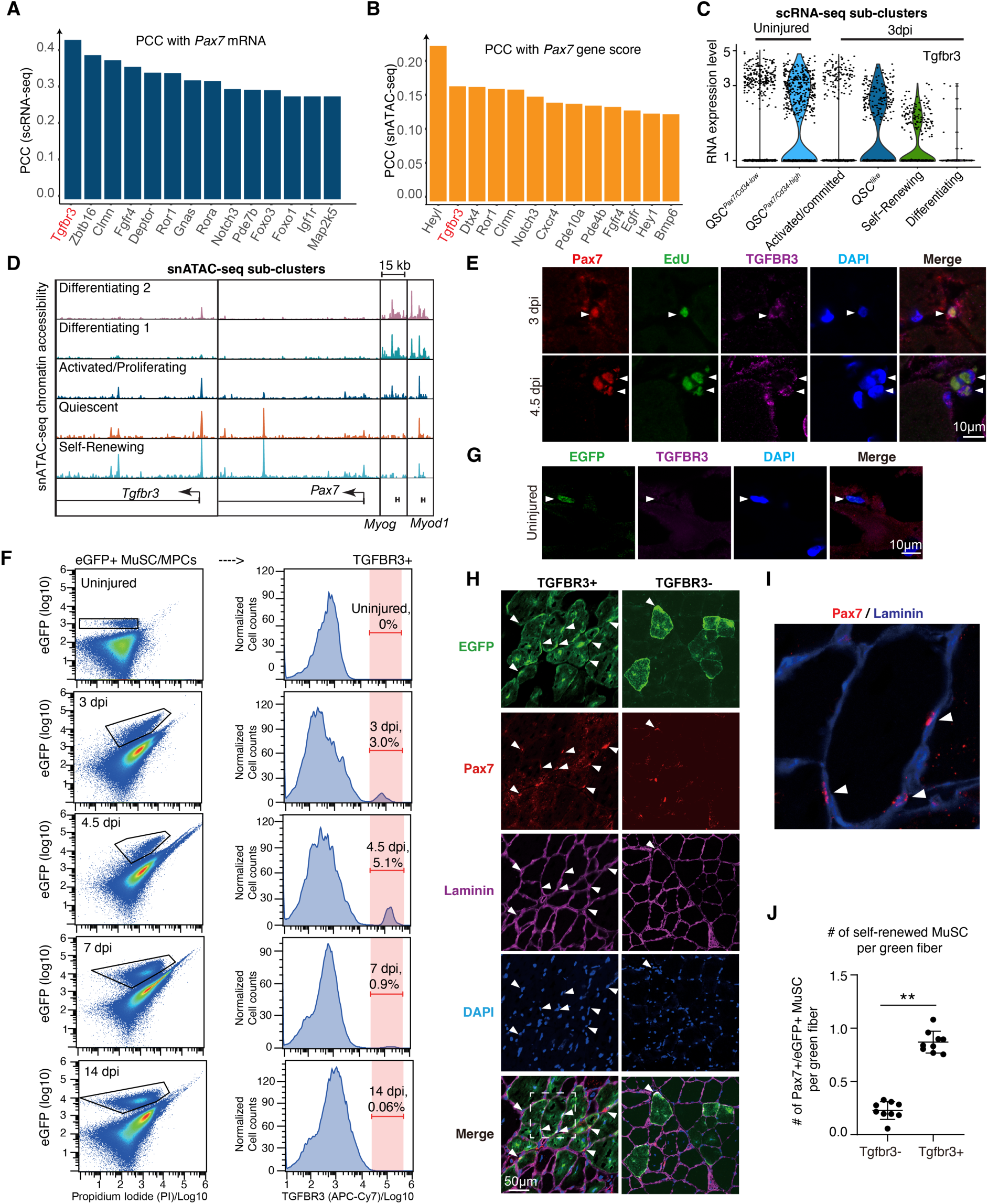
TGFBR3 protein expression identifies self-renewing MuSCs that contribute to the *in vivo* stem cell pool in muscle regeneration. (**A, B**) Pairwise Pearson correlation coefficient (PCC) was calculated using (**A**) mRNA and (**B**) gene score of all the genes against that of Pax7 across all the single cells of MuSC/MPCs. The top-ranked genes with the most robust and significant PCC with Pax7 were plotted. **(C)** Violin plot showing Tgfbr3 mRNA expression across the six sub-clusters of MuSC/MPCs defined in Fig 2C. **(D)** Genome browser view of chromatin accessibility at the indicated gene loci of *Tgfbr3*, *Pax7*, *Myod1* and *Myog* within the five sub-clusters of MuSC/MPC defined by snATAC-seq shown in Fig 2A. The height of each tracks normalized against sequencing depth. **(E)** Acute muscle injury was induced in TA muscle by intramuscular injection of BaCl_2_. At 3dpi (top) or 4.5dpi (bottom), EdU solution was injected intraperitoneally to label the proliferating cells in vivo. TA muscle was collected at four hours post EdU injection for cross-sections, followed by immuno-fluorescence staining. Arrowhead: the Pax7+/EdU+/TGFBR3+ cells, N = 3. **(F)** The lower hindlimb muscle of Pax7/eGFP mice were used to prepare the single-cell suspensions for FACS analysis. Cells were prepared from uninjured musle (top row), and the muscle at 3 days post injury (dpi) (second row), 4.5dpi (third row), 7 dpi (4th row) and 14 dpi (5th row), respectively. The cells were stained with APC-Cy7-conjugated TGFBR3 antibody prior to FACS. Pink bar highlights the fraction of TGFBR3+ cells (right panels) within the eGFP+ populations (left panels). **(G)** The TA muscle of uninjured Pax7/eGFP reporter mice were collected for immunostaining analysis. N = 3 **(H, I, J)** Following the same FACS condition described in (**F**), using 3dpi mice, the eGFP+/TGFBR3+ and the eGFP+/TGFBR3-cells were freshly sorted separately. The equal amount of eGFP+ cells were transplanted into TA muscle of recipient mice (10,000 cells per TA) that do not express eGFP transgene. The TA muscle of recipient mice were pre-injured one-day prior to the transplantation. 30 days post transplantation, the TA muscle was collected for the analysis. (**H** and **I**) The immunostaining of TA muscle cross-sections. Arrowheads point to the eGFP+/Pax7+ mononuclear MuSCs surrounding newly formed eGFP+ myofibers. **(I)** A close-up view of the dash-line boxed region in (**H**, bottom row, TGFBR3+) for further visualization of PAX7 and Laminin staining on TA muscle cross-section. **(J)** Quantification of eGFP+ MuSCs on twenty cross-section slides of TA muscle collected from five mice. **: t-test P < 0.01. N = 3.

To visually assess the expression of TGFBR3 protein in MuSC/MPCs in regenerating muscle *in situ*, we focused on 3 and 4.5dpi muscle, which is about the times we observed the most abundant self-renewing MuSCs from our snATAC-seq analyses (**Fig 2D**). We injected 5-Ethynyl-2’-deoxyuridine (EdU) intraperitoneally (IP) to label cycling cells in vivo 4 hours before tissue collection. At both 3dpi and 4.5dpi, we detected PAX7+ and EdU+ double-positive MuSCs expressing TGFBR3 protein (**Fig 4E**, arrowhead). Particularly, at 4.5dpi, we found that many MuSC/MPC clusters comprise Pax7+/EdU+ double-positive cells, suggesting that they are clonally expanded MuSC/MPCs.

To further assess the temporal dynamics of TGFBR3 expression in MuSC/MPC during muscle regeneration, we permanently marked MuSC/MPCs with a Pax7-CreER-based tagging strategy. Two weeks after tamoxifen injection (once per day for a total of 5 consecutive days), we induced acute injury of TA muscle by BaCl_2_ injection, followed by FACS analysis of TGFBR3 expression in the eGFP+ MuSC/MPCs before and after injury at days 3, 4.5, 7, and 14 (**Fig 4F**). We found that the TGFBR+ MuSC/MPCs account for ∼3.0%, 5.1%, and 0.9% of eGFP+ cells at 3, 4.5, and 7 dpi, respectively. In the fully repaired muscle at 14dpi, we detected very few eGFP+ cells expressing TGFBR3+ (0.06%) (**Fig 4F**). By contrast, the eGFP-labeled quiescent MuSCs in uninjured muscle did not express measurable TGFBR3 protein, as indicated by both FACS analysis (**Fig 4F,** top row) and immunostaining (**Fig 4G**). These results indicate that TGFBR3 protein is preferentially expressed in MuSC/MPCs that are undergoing self-renewing process during active regeneration, but not in those either in quiescent or have completed the process. 2.TGFBR3 is not measurably present in quiescent MuSCs, even though they express *Tgfbr3* transcripts (**Fig 4C**).

Cell transplantation is the gold standard to assess the self-renewal capacity of MuSC/MPCs *in vivo*. To determine whether the TGFBR3+/eGFP+ double-positive MuSCs are the bonafide self-renewing MuSCs, we followed an established protocol to transplant the freshly isolated MuSCs into mice for in vivo lineage tracing in regenerating muscle.^12^ About 10,000 TGFBR3+/eGFP+ double-positive cells were freshly sorted from 3dpi muscle (**Fig 4F,** 3dpi), and the same number of TGFBR3-/eGFP+ cells were collected as control. We induced acute TA muscle injury by injecting BaCl_2_ into the recipient mice that do not express eGFP transgene. The FACS-sorted eGFP+ cells were injected into the injury site one day later. At 30dpi, the fully repaired TA muscles were collected for analysis of eGFP+ satellite cells and muscle fibers. At the cross-section of TA muscle, we found that both TGFBR3+ and TGFBR3-cells similarly contributed eGFP+ myofibers (**Fig 4H**). However, compared to the TGFBR3-cells, the TGFBR3+ cells were ∼4.8-fold more efficient to repopulate the Pax7+ cells surrounding the eGFP+ myofiber *in vivo* (**Fig 4H, 4I**, arrowheads; **Fig 4J,** quantification). Notably, these graft-derived, eGFP+ mononuclear cells express PAX7, and are located in the periphery of the myofiber inside the basal lamina, the known anatomical position of self-renewed MuSC (**Fig 4I**).

Previous studies showed that the freshly isolated, uninjured quiescent MuSCs can potently repopulate the in vivo MuSC pool ^11–18^. Because the uninjured quiescent MuSCs do not express TGFBR3 protein, in our transplantation experiments, we concluded that the transplanted TGFBR3+/eGFP+ cells had very few if any quiescent MuSCs that may remain due to incomplete injury. Therefore, the self-renewed eGFP+ MuSCs and the green myofibers observed in the repaired muscle are derived from the TGFBR3+/eGFP+ self-renewing MuSCs, and do not represent the remaining quiescent MuSCs due to incomplete injury. Taking all of our results into consideration, we conclude that TGFBR3 is a unique marker of the self-renewing sate of MuSCs, allowing their isolation and transplantation to injured muscle. Importantly, engrafted, TGFBR3+ self-renewing MuSCs potently contribute to new myofibers as well as repopulate the stem cell niche.

### SMAD4 is a crucial transcription factor required for MuSCs self-renewal in vivo

As a co-receptor in the TGFβ superfamily, TGFBR3 possesses no kinase activity on its own. Instead, it can either potentiate or inhibit BMP and TGFβ pathways by interacting with distinct ligands and receptors, and/or acting through its ectodomain shedding process ^40, 41^. The transcription factors Smad2/3 and Smad1/5 are Receptor-regulated Smads (R-Smads) that lie downstream of TGFβ and BMP signaling pathways, respectively ^42^. Because TGFBR3 marks self-renewing MuSCs in our experiments, we hypothesized that SMAD proteins are involved in the cell fate decisions of MuSCs. Consistent with this idea, from our snATAC-seq data, we noticed changes in related sequences acquiring open chromatin structure in MuSC/MPCs undergoing both self-renewal and differentiation fates (**Fig 5A**). Following the self-renewal trajectory defined by snATAC-seq analysis, the Smad2/3 binding motif is enriched in sequences acquiring open chromatin during regeneration, and then depleted from such regions as quiescent MuSCs transition to the self-renewal state (**Fig 5A**, top left). Interestingly, the enrichment of Smad1/5 motifs in the open chromatin sequences follow a complementary pattern: it is gradually decreased in accessible regions of quiescent MuSCs as regeneration progresses, then slightly increased in self-renewing cells (**Fig 5A**, bottom left). Similarly, we also observed dynamic and complementary changes of enrichment of binding motifs of Smad2/3 and Smad1/5 in the DNA sequences acquiring accessibility in the MuSCs/MPCs following the differentiation trajectory (**Fig 5A**, right). In addition to these chromatin accessibility dynamics, transcriptomic evidence also suggests the differential regulation of TGFβ superfamily signalings (**Fig 5B**, **Table S6**, Gene Ontology analysis), as well as the R-Smads and their transcriptional targets **(Table S5, Fig S2C - S2G)** in SRSCs compared to the activated/committed MPCs and quiescent MuSCs. Together, these results strongly support the idea that the fate choice of MuSCs to undergo self-renewal versus commitment to differentiation during regeneration is partly regulated by the Smad transcription factors.

**Figure 5.**
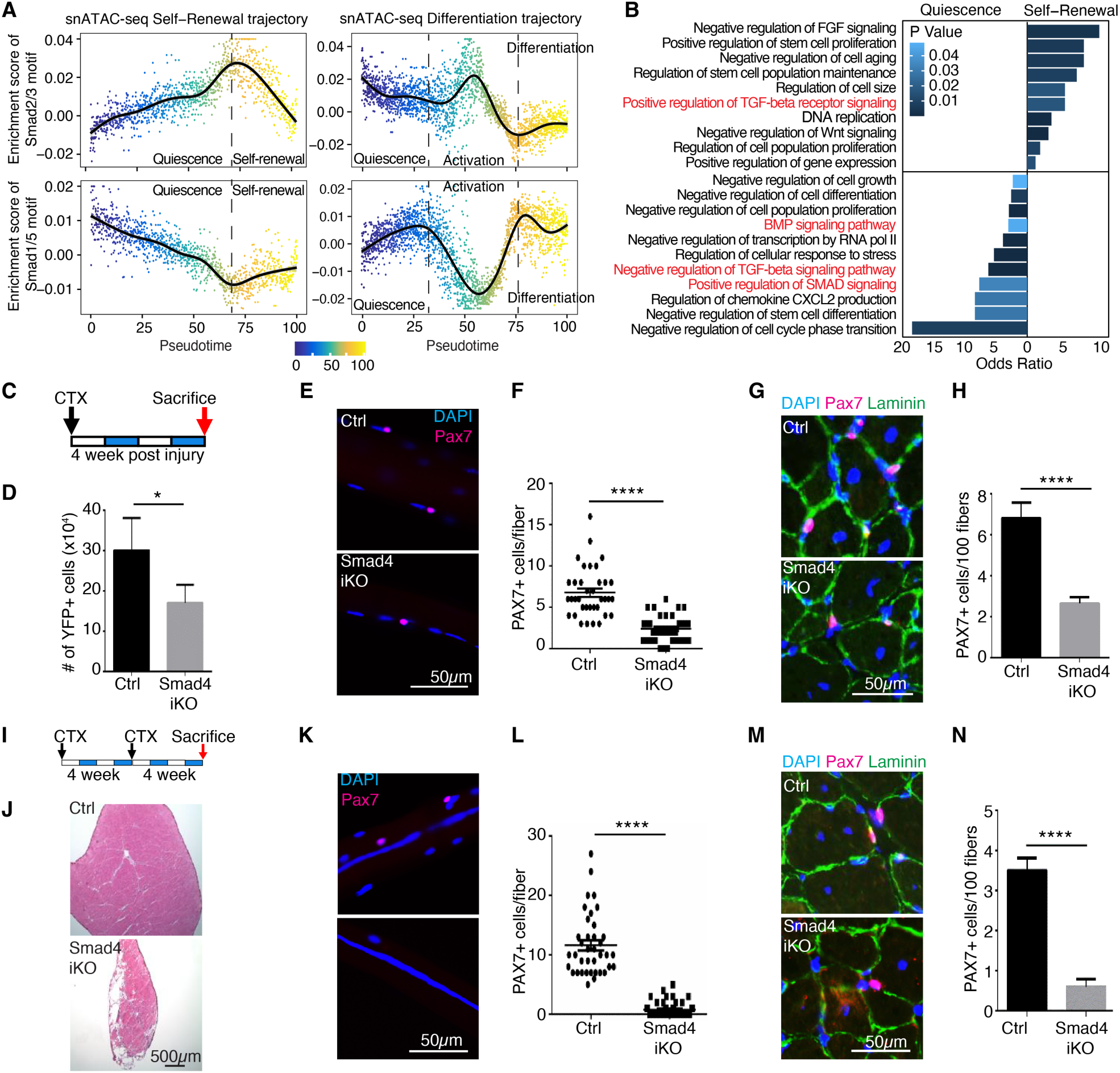
SMAD4 is a crucial transcription factor required for MuSC self-renewal in vivo during regeneration. **(A)** Scatter plot showing the enrichment score (y-axis) of the binding motifs for Smad2/3 (top), and Smad 1/5 (bottom) of each single cell aligned along the MuSC self-renewal trajectory (left, x-axis) and the myogenic differentiation trajectory (right, x-axis). Both trajectories are defined by snATAC-seq analysis shown in Fig 2F**, 2G**. Color key: pseudotime. The dashed line separates the Quiescent, Self-renewing, activation, and differentiation sub-clusters. **(B)** The significantly enriched gene ontology (GO) terms (p-value < 0.05) were identified using the differentially expressed genes in the genuine quiescent versus self-renewing MuSCs defined by scRNA-seq in Fig 3C. Differentially expressed genes: FDR<0.05 (Wilcoxon rank-sum test) and absolute Log2 (fold change) > 0.6. **(C)** Schematic of the experimental design for self-renewal assay. Hindlimb muscles of Pax7/YFP mice, after tamoxifen administration (descripted in **Fig S3B**), were injured by cardiotoxin (CTX) injection and collected after four weeks for further experiments described in (**D-H**). **(D)** 4-week after CTX injury, MuSCs were sorted from Ctrl and Smad4 iKO mice and the cell numbers were recorded. **(E, F)** 4-week after CTX injury, EDL fibers (n>30) were isolated from Ctrl and Smad4 iKO mice, and subjected to immunostaining for Pax7. The nuclei were counterstained with DAPI. The images are representative of multiple fibers. Scale bars, 50 µm. **(F)** Quantification of Pax7+ MuSC number from (**E**). **(G, H)** 4-week after CTX injury, TA cross-sections of Ctrl and Smad4 iKO mice were subjected to immunostaining for Pax7 and Laminin. The nuclei were counterstained with DAPI. The images are representative of multiple sections. Scale bars, 50 µm. **(H)** Quantification of Pax7+ cell number per 100 myofibers on TA sections in (**G**). **(I)** Schematic of the experimental design for self-renewal assay using repeated injury. After tamoxifen administration (descripted in Fig S3B), the lower hindlimb muscles of Pax7/YFP mice were injured by CTX twice with 4-weeks interval. Mice were sacrificed 4 weeks after the second CTX injury, hindlimb muscles were collected and subjected to further assays described in (**J-N**). **(J)** 4-week after the second CTX injury, TA cross-sections were subjected to H&E staining. The images are representative of multiple TA cross-sections from three pairs of littermates. Scale bars, 50 µm. **(K, L)** EDL fibers (n>30) were isolated from the same mice in (J) and subjected to immunostaining for Pax7. The nuclei were counterstained with DAPI. The images are representative of multiple fibers. Scale bars, 50 µm. **(L)** Quantification of Pax7+ MuSC number in (K). **(M, N)** Twice-injured TA cross-sections were subjected to immunostaining for Pax7 and Laminin. The nuclei were counterstained with DAPI. The images are representative of multiple sections. Scale bars, 50 µm. **(N)** Quantification of Pax7+ cell number per 100 myofibers on TA cross-sections in (M). All the results from **D** to **N** are presented as mean ± SEM. *p < 0.05, **p < 0.01, ***p < 0.001, ****p < 0.0001. Numbers of biological replicates are: N=4 in (**D**) and N=3 in (**E-N**).

To test this hypothesis, we disrupted *Smad4*, the Co-Smad required for the activity of all the R-Smads (Smad2/3 and Smad1/5/8), using mice with Loxp-flanked Smad4 exons. We crossed *Smad4^flox/flox^* (“f/f” thereafter) mice with *Pax7^CreER/CreER^: Rosa26^YFP/YFP^* (*Pax7/YFP*) mice (i.e., *Smad4 iKO*), enabling use of YFP as a reporter to mark MuSCs (**Fig S3A-S3C**). The KO efficiency of SMAD4 in MuSCs is validated by Western blot at the protein level (**Fig S3D**, top) and by RNA-seq at the RNA level as indicated by lacking of transcripts mapped to *Smad4* exon 8 (**Fig S3D**, bottom). Four weeks after acute muscle injury (**Fig 5C**), we noticed a 50% reduction in the number of YFP+ MuSCs in *Smad4* iKO samples compared to that from heterozygous mice (**Fig 5D**). A similar decrease was detected by quantification of self-renewed MuSCs in isolated myofibers (**Fig 5E, 5F**) and TA muscle cross-sections (**Fig 5G, 5H**). The observed reduction of MuSCs in the fully repaired muscle suggest self-renewal defect of SMAD4 mutant MuSCs. Given these observations and based on findings by others ^28, 43^, we hypothesized that a second injury might induce a more severe regeneration defect and even fewer self-renewed MuSCs. Indeed, following a sequential injury scheme (**Fig 5I**), we found that TA muscle regeneration in *Smad4* iKO mice was much less efficient than in heterozygous mice (**Fig 5J**). Notably, the freshly isolated single myofibers or TA muscle cross-sections from *Smad4* iKO mice had few if any detectable Pax7+ cells (**Fig 5K, 5L** for myofiber and **Fig 5M, 5N for TA sections**). Taken together, we conclude that *Smad4* is essential in adult MuSCs for satellite stem cell self-renewal during regeneration.

### Mechanisms of control of muscle stem cell self-renewal by *Smad4*

To identify potential mechanisms by which Smad4 regulates MuSCs, we carried out a series of analysis of Smad4 iKO mice. We found that SMAD4 ablation in adult MuSCs does not measurably affect stem cell maintenance (**Fig S3D-3J**), exit from quiescence and cell cycle re-entry (**Fig 6A, 6B**), or the early phases of cell proliferation (**Fig 6C, 6D**). Interestingly, we found that SMAD4 is required for prolonged proliferation of MPCs (**Fig 6C, 6D**). Therefore, we hypothesized that SMAD4 plays a critical role in the early activated and proliferating MuSC/MPCs by controlling the balance between self-renewal versus myogenic differentiation fate.

**Figure 6.**
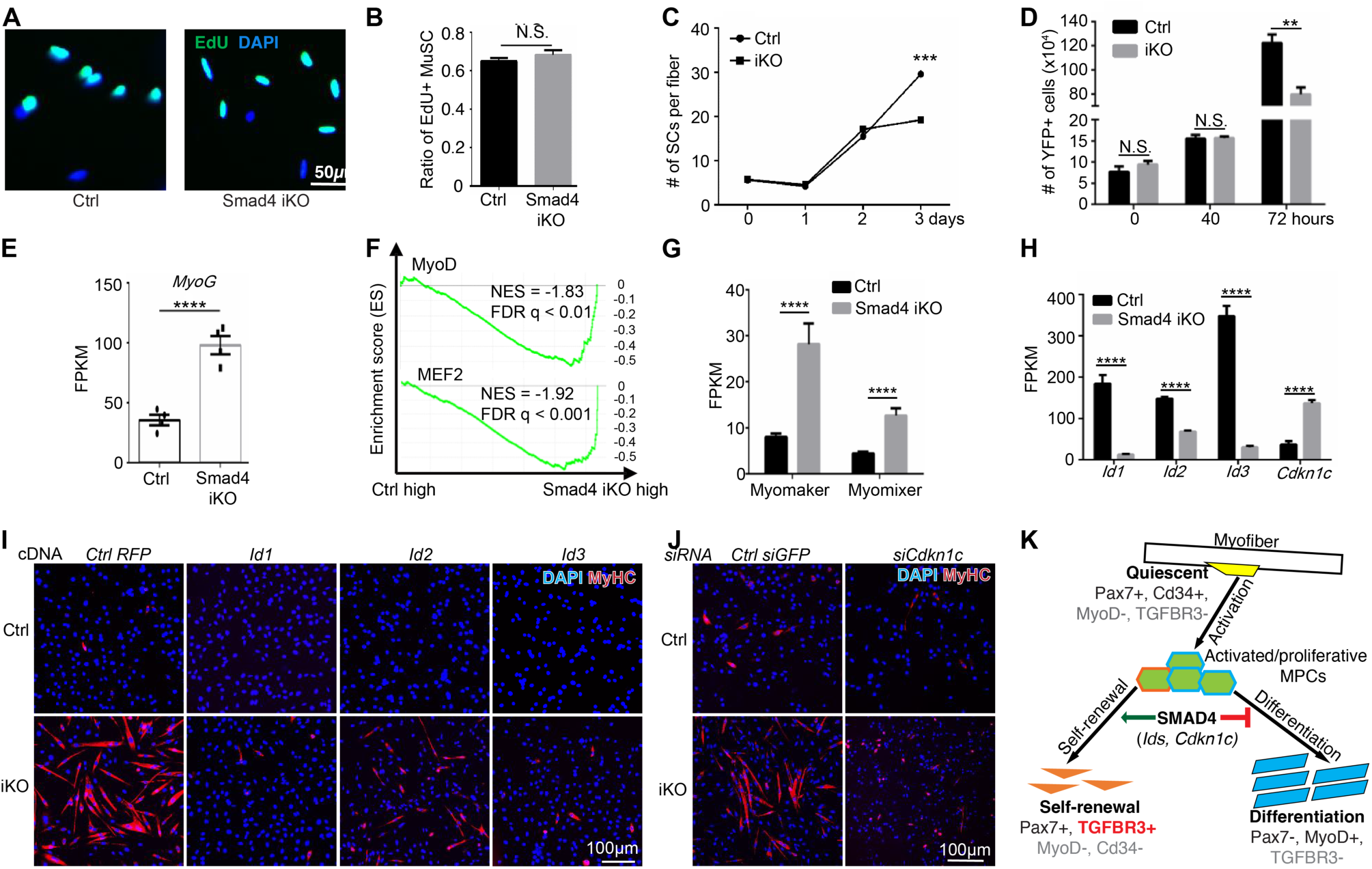
SMAD4 promotes self-renewal by restricting the alternative differentiation fate of MuSCs. (**A, B**) Equal number of YFP+ MuSC were FACS-sorted from Ctrl versus SMAD4 iKO mice, and cultured in the growth media supplemented with EdU. After 40 hours of in vitro culture, the cells were subjected to staining for EdU. Nuclei were counterstained with DAPI. Scale bars, 50 µm. **(B)** Quantification of the percentage of EdU+ cells in (**A**). **(C**) Myofibers were isolated from EDL muscle of Ctrl and Smad4 iKO mice, respectively. The myofiber were cultured for 0-, 1-, 2-, and 3-days before fixation. The number of YFP+ cells per fiber was quantified and shown. **(D)** The YFP+ MuSCs were isolated from lower hindlimb muscles of Ctrl and Smad4 iKO mice respectively at indicated times post CTX injury. The numbers of MuSCs were recorded by FACS. **(E-H)** 40-hour post CTX induced injury, the YFP+ MuSC/MPCs were FACS sorted from the injured lower hindlimb muscle of control and SMAD4 iKO mice. The collected cells were subjected to RNA-seq analysis. N = 4 **(E, G, H)** Bar plots represent the normalized reads counts (mean FPKM values) for (**E**) *Myogenin;* (**G**) *Myomaker* and *Myomixer;* and (**H**) *Id1, Id2, Id3* and *Cdkn1c* in MuSCs from 4 pairs of littermates. Error bars were calculated by Cuffdiff 2. **(F)** Gene Set Enrichment Analysis results showing MyoD and MEF2 target genes are significantly enriched in SMAD4 iKO samples. **(I, J)** Equal number of FACS-isolated uninjured MuSCs were cultured for 18h before: (**I**) treatment of adenovirus to overexpress RFP, Id1, Id2, and Id3 cDNA for 6h; or (**J**) transfection of siRNAs against control *GFP* and *Cdkn1c*. After another 36h, the cells were fixed and subjected to immunostaining for MyHC (red). The nuclei were counterstained with DAPI (blue). Scale bars, 100 µm. **(K)** Schematic illustration showing the expression of the indicated genes at the protein levels at different states of MuSCs/MPCs during muscle regeneration. After injury, the quiescent MuSCs (Pax7+, Cd34+, MyoD1-, TGFBR3-) gave rise to a heterogeneous population of activated and proliferative MuSC/MPCs. SMAD4 plays a critical role in regulating the cell fate decision of these activated/proliferative MPCs by targeting *Ids* and *Cdkn1c* genes. Specifically, SMAD4 restricts the precocious differentiation fate and promotes self-renewal, which is marked by TGFBR3 protein expression. All the results are presented as mean ± SEM. *p < 0.05, **p < 0.01, ***p < 0.001, ****p < 0.0001. Numbers of biological replicates are: N=3 in (**A-D**), N=4 in (**E-H**), and N=3 in (**I-J**).

To test this idea, we assessed sorted MuSC/MPCs from hindlimbs of control and SMAD4 iKO mice by RNA-seq analysis at 40hpi, the time when the activated MuSC/MPCs are poised to enter their first cell cycle for either self-renewal or differentiation (**Fig 2D**). Unexpectedly, at 40hpi, the *Myog* mRNA levels in *Smad4*-null MuSCs were ∼2.8 times higher than those in control samples (**Fig 6E**). The GSEA analysis revealed that the myogenic commitment/differentiation-related genes, including *MyoD1* and *Mef2a* target genes, were significantly enriched in the differentially expressed genes in *Smad4*-null MPCs compared to the wild-type control MPCs (**Fig 6F**). Moreover, *Myomaker* and *Myomixer*, two genes essential for myoblast fusion and expressed in the terminally differentiated myocytes ^44, 45^, were also present at higher levels in *Smad4*-null MPCs at 40hpi (**Fig 6G**). These results supported the notion that *Smad4-null* MuSCs prematurely commit to myogenic differentiation as early as 40 hpi, a timepoint during which the control activated MuSCs are largely proliferating.

In *Smad4*-null MuSCs, we also found that the mRNA levels of the *Id* family genes are lower than those of controls, whereas *Cdkn1c* (*p57Kip1*) levels are higher (**Fig 6H**). Ids are well-characterized negative regulators for myogenic differentiation, as they prevent MyoD1/Myf5 from binding to DNA by complexing with either MyoD1/Myf5 or E2A proteins ^46^. By contrast, p57Kip1, a cell cycle inhibitor, promotes cell cycle exit at the onset of myogenic differentiation. Strikingly, we found that the re-expression of individual members of *Id* genes (**Fig 6I**), or knockdown of *p57Kip1* (**Fig 6J**), effectively prevented precocious differentiation of in primary cultures of freshly isolated *Smad4*-null MPCs. These results indicate that *Id* family genes and *Cdkn1c* are key downstream targets of Smad4 that function to prevent precocious myogenic differentiation of activated/proliferative MuSCs.

In summary, we propose that Smad4 is key to control the balance between self-renewal versus differentiation fate of activated MuSCs at early timepoint of regeneration. Mechanistically, *Ids* and *p57Kip1,* the downstream target genes regulated by SMAD4, play a critical role in restricting the onset of a precocious myogenic differentiation process. This crucial mechanism enables a subset of activated/proliferative MPCs to enter the self-renewal state marked by TGFBR3 (**Fig 6K**).

## DISCUSSION

The ability to leverage tissue-resident stem cells for future regenerative therapeutic strategies is currently hampered by the significant gap in our knowledge of the identity and mechanisms of stem cell self-renewal. In skeletal muscle, the attempt of such a therapeutic approach can be traced to 1978 when Partridge and colleagues demonstrated the fusion between host and donor myoblasts in skeletal muscle grafts ^47^. However, as soon as primary MuSC are isolated from their in vivo niche, they immediately lose their stem cell potency and are committed to the differentiation fate ^11–18^. Such committed MPCs can contribute to new myofibers, but are not efficient to repopulate the in vivo stem cell pool, nor to support long-term muscle homeostasis and regeneration ^11^. The existence of self-renewing population of MuSCs in vivo was already predicted, but the in vivo self-renewing state of MuSCs has not been characterized. Here, through single-cell chromatin accessibility profiling, we uncover the molecular identity of a population of self-renewing MuSCs as well as a transcriptional program that governs their regeneration potential. Since the self-renewing MuSC/MPCs have a higher efficiency of engraftment than most proliferating MPCs, this finding is of potential practical importance for transplantation therapeutics. The capacity of identifying and isolating the self-renewing MuSCs from the regenerating muscle will allow for future research opportunities to investigate the mechanism of muscle regeneration.

Our results indicate that SMAD4 is a crucial transcription factor controlling the cell fate choice of early activated/proliferative MuSC for self-renewal versus differentiation. Among the eight known Smad proteins in mammals, *Smad4* is unique in that it functions as an indispensable Co-Smad for five R-Smads (receptor-regulated Smads, i.e., *Smad1, 2, 3, 5*, and *8*): upon activation of their receptors by members of the TGFβ superfamily, Smad4 forms heterodimers with one of the five R-Smads in the cytosol and together they are translocated to the nucleus to bind to their target genes ^48^. The function of SMADs, including SMAD4, has been studied in the processes of myoblast proliferation and differentiation ^49–54^. Nevertheless, whether and how the SMADs proteins causally regulates MuSC self-renewal was unknown. Recently, Chakkalakal and his colleagues induced Smad4 ablation in adult MuSCs and showed that the injury-induced muscle regeneration was compromised due to precocious differentiation of the mutant MuSCs ^53^. Our study further demonstrates the SMAD4 not only plays a critical role in regulating myogenic differentiation, but also serves as a key regulator for MuSC self-renewal in regeneration.

It should be noted that the functional significance of the elevated expression of TGFBR3 protein in the self-renewing MuSCs remains to be elucidated. As a co-receptor of TGFβ superfamily, TGFBR3 can serve to either activate or silence BMP or TGFβ signaling ^40, 41^. Intriguingly, the expression of TGFBR3 protein coincides with the decreased SMAD2/3 activity, indicated by less enriched binding motif of SMAD2/3 in the open chromatin sequences of the self-renewing population. Although activation of SMAD2/3 via TGFβ or myostatin signaling inhibits myogenesis ^55, 56^, recent studies reported that SMAD2 and SMAD3 can promote myogenic differentiation independent of TGFβ signaling ^52, 57–59^. Particularly, SMAD2 is identified as an important positive regulator of myogenic differentiation by driving *Myog* expression ^52^. These results suggested that in the early activated MuSCs engaged in self-renewal, the inactivation of SMAD2/3 could potentially block the myogenic differentiation fate to enable self-renewal. In the future study, it would be very interesting to employ the sophisticated genetically engineered mouse models to determine whether TGFBR3, SMAD2/3 (double knock-out), and SMAD1/5/8 (triple knock-out) play a causal role in controlling MuSC fate choice.

In conclusion, with time-resolved single-cell atlas of chromatin accessibility of skeletal muscle regeneration, we have delineated the self-renewal versus myogenic differentiation trajectories of MuSC upon injury. Our results identify TGFBR3 protein as a unique marker of self-renewing MuSCs, and uncover the critical role of SMAD4 for MuSC self-renewal and in controlling the fate choice of early activated/proliferative MuSCs.

## ACCESSION CODES AND DATA AVAILABILITY

Sequencing data have been deposited to the NCBI Gene Expression Omnibus (GEO) (http://www.ncbi.nlm.nih.gov/geo) under accession number GSE199499. The publicly available single-cell RNA-seq data used have the accession numbers SRR10870296, SRR10870297, and SRR10870298. Additional materials, data, code, and associated protocols are available upon request.

## ACKNOWLEDGEMENTS

We thank Drs. Brigid Hogan (Duke University) and Kenneth Poss (Duke University) for feedback on previous versions of the manuscript, and Dr. Sharyn Endow (Duke University) for helping Tn5 purification used in our snATAC-seq experiments. This work is supported by a new lab startup fund from Duke University Regeneration Next Initiative (current Duke Regeneration Center) (to Y.D.), Duke Whitehead Scholarship (to Y.D.), Glenn Foundation for Medical Research and AFAR Grants for Junior Faculty (to Y.D.), and NIH 4D Nucleome Consortium U01HL156064 (to Y.D.).

## AUTHOR CONTRIBUTIONS

X.L., C.S., Xiaolin W. and Xiuqing W. performed the experiments. A.E.O. led the effort of sequencing data analysis with help from Y.X. A.E.O., X.L., C.S., Z.W., and Y.D. wrote the paper.

## MATERIALS AND METHODS

### Animals

All wild-type (WT) and transgenic animals were on the C57BL6 background strain and were between 3-6 months of age. Pax7CreERT2/+ (Strain #: 017763), NuTRAP/+ (Strain #: 029899) and Rosa26-YFP (Strain #: 006148) mice were purchased from Jackson Laboratory, and Smad4 flox/flox mice were kindly provided by Prof. Huiyao LAN (The Chinese University of Hong Kong), and Pax7CreERT2/+; NuTRAP/+, Pax7CreERT2/+; ROSA26 YFP/+ and Pax7CreERT2/+; ROSA26YFP/+; Smad4 flox/flox were generated by crossing Pax7CreERT2/+ mice with NuTRAP/+ or ROSA26YFP/+ mice or Smad4 flox/flox crossed with Pax7CreERT2/+; ROSA26YFP/+. Tamoxifen is used to activate Cre in mice. Dissolve tamoxifen (Sigma-Aldrich, T5648) in corn oil at a concentration of 20 mg/ml and mice received approximately 75 mg tamoxifen/kg body weight via intraperitoneal injection for 5 consecutive days. All the experiments with animals were approved by the Institutional Animal Care and Use Committee (IACUC) at Duke University or the Animal Ethics Committee at Hong Kong University of Science & Technology (HKUST).

### Muscle injury using BaCl^2^ and CTX

To activate MuSC in vivo, 1.2% BaCl_2_ (w/v in H2O) or 10 μM Cardiotoxin (CTX) was used to induce acute injury. Mice were injured by injecting 30 μL of BaCl2 or CTX to Tibialis anterior (TA) muscles for immunofluorescence staining or 50 μL of BaCl2 or CTX into the lower hind-limb muscles (below the knee) for flow cytometry to isolate single-nucleus suspension, single-cell suspension and muscle satellite cells(MuSC).

### Single Nucleus ATACseq experiment

Injury and no injury lower hind-limb muscles were snap-frozen and sectioned on dry ice, and then homogenized on a gentleMACS Octo Dissociator (Miltenyi) using the “Protein_01_01” protocol with MACS buffer (5 mM CaCl2, 2 mM EDTA, 1X protease inhibitor (Roche, 05-892-970-001), 3 mM MgAc, 10 mM Tris-HCL pH 8, 0.6 mM DTT) and went through 30 μM CellTrics filter (Sysmex, 04-004-2326) into 15 mL tubes. After 500 x g, 5 min, 4°C centrifuge, the pellet was resuspended in 3 mL Nuclear Permeabilization Buffer (1X PBS, 5% Bovine Serum Albumin, 0.2% IGEPAL CA-630 (Sigma), 1 mM DTT, 1X Protease inhibitor) and were rotated at 4°C for 5 minutes. Centrifuge and remove the supernatant, the permeabilized nuclei were resuspended in 500 μL high salt tagmentation buffer (36.3 mM Tris-acetate (pH = 7.8), 72.6 mM potassium-acetate, 11 mM Mg-acetate, 17.6% DMF) and counted using a hemocytometer. Concentration was adjusted to 1,000 nuclei/9 μl, and 1,000 nuclei were dispensed all four 96-well plates (in total of 384 wells). For tagmentation, 1 μL barcoded Tn5 transposomes (**Table S7**) were added using a BenchSmart 96 (Mettler Toledo), and incubated for 90 min at 37°C with shaking (900 rpm). To inhibit the Tn5 reaction, 2.5 μL of 100 mM EDTA (final 20mM) were added to each well with a BenchSmart 96 and the plate was incubated at 37°C for 15 min with shaking (900 rpm). Next, 6 μL of 3x sort buffer (3% BSA, 3 mM EDTA in PBS) were added using a BenchSmart 96. All 384 wells were combined into a separate FACS tube and stained with Draq7 at 1:500 dilution (Cell Signaling). We used a Sony SH800 Sorter (Sony) to sort each well containing 80 nuclei per well into new four 96-well plates (in total of 384 wells) containing 8 μL primer dilution buffers (15 pmol primer i7, 15 pmol primer i5, 200 ng BSA (Sigma), 0.027% SDS, (**Table S7**). The 96 well plate was incubated at 65°C for 25 min in the PCR and 4 μL 1.33% Triton-X was added to each well to quench the SDS. Subsequently, 11 μL Q5 mix (5 μL 5 X Q5 buffer, 0.5 μL 10 mM dNTP, 0.25 μL Q5 polymerase, 5.25 μL H2O) were added to each well and samples were PCR-amplified (72°C 4 min, 98°C 30 s, (98°C 10 s, 60°C 30 s, 72°C 45 s) × 12 cycles, held at 4°C). After PCR amplification, all wells were pooled to purify DNA products using the Zymo DNA Clean & Concentrator-5 kit (Zymo, Cat #: 11-303) according to manufacturer’s protocols. Purify DNA products was ran in the 1.5% Argo gel and 200∼600 bp DNA library was collected, and the resulting DNA libraries were purified according to the Zymo clean Gel DNA Recovery Kit (Zymo, Cat #: 11-300) according to manufacturer’s protocols. Final libraries were quantified using a Qubit™ 1X dsDNA HS Assay Kit (Invitrogen, Q33231) and a nucleosomal pattern of fragment size distribution was verified using a High Sensitivity D1000 Tapestation (Agilent, 5067-5582).

### FACS sorting of single-cell suspension for MuSC

Isolation of single cell and MuSC by physical dissociation and enzymatic digestion was followed as previously described (Liu et al., 2015). In brief, we dissected the hindlimb muscles from the mouse and digested the minced muscles with collagenase II (800U/mL, (Worthington, LS004177)) in wash medium (F10, 10% horse serum, 1% P/S) for 90 mins. The tissue was further digested by collagenase II (100U/mL) and dispase (1.1 U/mL) in wash medium for 30 mins and passed ten times through a 20-gauge needle, resuspend with wash medium, and went through a 40 μm filter to get single cell suspension for cell sorting. Single cell suspensions from WT mice were stained with Propidium Iodide (PI, 1:1000), and then was sorted on a Sony SH800 sorter equipped with 405-nm, 488-nm, 561-nm, and 633-nm lasers. PI-single cell from WT mice was sorted to Single-cell RNAseq. GPF+; PI- or YFP+; PI-single cell from reporter mice was used to isolate MuSC. For MuSCs transplantation, single cell suspension was stained with TGFBR3 antibody (1: 200, Invitrogen, MA5-17187), and stained with APC/Cy7 Goat anti-mouse IgG (1:500, BioLegend, 405316). APC/Cy7 Goat anti-mouse IgG Purified Mouse IgG2b, κ Isotype Ctrl Antibody (1:500, BioLegend, 400302) or secondary single-stained was used as control, the TGFBR3+; GPF+; PI- and TGFBR3-; GPF+; PI-were sorted to transplant, respectively.

### Single Cell RNA-seq experiment

16k cells were counted using a hemocytometer, and loaded onto a 10x Genomics Chromium chip per manufacturer’s recommendations. Reverse transcription and library preparation was performed using the Chromium Next GEM Single Cell 3ʹ Reagent Kits v3.1 (10x Gemomis, CG000204) according to manufacturer’s protocols. After a cleanup with SPRIselect Reagent Kit (Beckman, C10640), final libraries were quantified using a Qubit™ 1X dsDNA HS Assay Kit (Invitrogen, Q33231) and fragment size distribution was verified using a High Sensitivity D1000 Tapestation (Agilent, 5067-5582).

### MuSC transplantation assay

After tamoxifen activating cre in Pax7creER/+;NuTRAP/+ mice, BaCl2 induced hind limb injury after 3 days, and freshly isolated MuSCs were collected by FACS sorting. Recipient TA muscles of C57BL6/WT mice were injured with BaCl2 and 24 hours in advance. Ten thousands of Tgfbr3+ and Tgfbr3-MuSC cells were injected into the injured left and right of recipient TA muscles. One month later, TA muscle will be collected for fixation and cryosection. Sectioned 10 μm slices were collected every 200 μm and subjected to immunostaining. All slices with GFP-positive cells around with GFP-positive fibers were regarded as a self-renewal MuSC and quantified and 3 maximum numbers in each TA muscle were used for statistical analysis.

### Isolation of single myofiber

Extensor digitorum longus (EDL) muscles were isolated and digested with Collagenase II (800 units/ml) in the wash medium at 37 ℃ for 75 min. Gentle trituration was performed to release single myofibers from the scattered EDL bundles in the wash medium. Healthy fibers were picked out and cultured in the wash medium for immunostaining.

### Bulk RNA-seq experiment

Four pairs of control and Smad4 iKO mice with the same age were injected with Tamoxifen to induce genetic recombination as described above. After CTX injured 40 h, MuSCs from the gaskin muscles were collected by FACS and then subjected to extract RNA with NucleoSpin® RNA XS kit from Macherey-Nagel. 1 ng high-quality RNA per sample were reverse transcribed and amplified into cDNA following Smart-seq2 library preparation protocol (Picelli et al., 2014). Thereafter, amplified cDNA was fragmented by sonication with a Covaris® S220 Focused-ultrasonicator. Nugen Ovation Ultralow Library System was the kit used for sample end repair, adaptor ligation, and amplification, in which 8 different barcodes were attributed to the 8 samples for identification. RNA and DNA qualities were verified with Agilent 2100 Electrophoresis Bioanalyzer Instrument. Finally, after preparation of a high-quality DNA library, pair-end RNA sequencing was done with Illumina Nextseq 500 system in Biosciences Central Research Facility in HKUST.

### Bulk RNA-seq data analysis

Upon acquiring the raw sequencing reads, the STAR aligner was applied to map the raw sequencing reads to the mouse genome assembly (mm10). 2-pass STAR alignment was conducted and then subjected to the gene differential expression analysis with Cuffdiff2, in which the significance cutoff was set at 0.05 (FDR-adjusted p_value). Aligned bam files were converted into bigwig files that could facilitate the visualization of sequencing data in the genome browser. Gene Set Enrichment Analysis (GSEA) was performed with the data set generated from the cuffdiff output.

### EdU labeling assay

In vitro, freshly sorted MuSCs were cultured in Ham’s F10 media with 10% horse serum (Corning, 35-030-CV) and constantly supplied with 10 mM EdU (Invitrogen, A10044) at the planned time points. For EdU incorporation assay in vivo, TA muscles of C57BL6/WT and Pax7creER/+; NuTRAP/+ mice were injured with BaCl2 after 3 or 4.5 days, and EdU was administered through intracellular injection at a concentration of 150 ug/g of body weight. Mice will be sacrificed after 4 hrs or 1 month, MuSCs were then fixed and TA muscles were collected to prepare cryosection, and stained using the Click-iT EdU Imaging Kit (Invitrogen, C10337) according to the manufacturer’s instructions.

### Histology and Immunostaining of muscle cross-section

TA muscles were embedded into Neg-50™ medium (Thermo, 6502) and put into liquid Nitrogen for freezing immediately after dissection. The TA muscles were sliced into 10 μm sections. For the hematoxylin-eosin staining, sections were stained with hematoxylin (Agilent, S3309) and eosin (Sigma, HT110216) and photographed by using a Leica DM5500B microscope (Leica). For immunostaining, cells and single fiber and sections were fixed with 1% PFA for 10 min and permeabilized (0.3% Triton X-100, 0.1M Glycine, PBS) for 30 min, and were blocked with Blocking buffer (4% BSC (Rockland, RLBSA50) and 1% goat serum (Gibco, PCN5000)). For Pax7 staining, sections were washed in PBS and then antigen-retrieved with sodium citrate buffer (10 mM, 0.05% Tween in PBS) at 95 ℃ for 30 min, and were washed and blocked with M.O.M. blocking reagent (Vector Laboratories, MKB-2213-1) according to the instruction. Incubations with primary antibodies (Pax7 (1:200, DSHB, AB_528428), Tgfbr3 (1:200, Sigma, SAB4502962), Laminin (1:500, Sigma, L9393), GFP (abcam, ab13970) and MyHC (1:200, DSHB, AB_528352)) were performed overnight at 4 ℃. Incubation with secondary antibodies conjugated to Alexa Fluor (Life Technologies) was performed for 1 hour at room temperature. For section, 0.1% sudan black was used to block the autofluorescence. After several washes with PBS, sections were stained with Mounting Medium With DAPI (abcam, ab104139). Immunofluorescence was performed using a Zeiss 880 upright confocal Airyscan (Zeiss) and Nikon fluorescent microscope (Ni-U). Imaging data acquisition was performed using ImageJ software.

### snATAC-seq data preprocessing

Raw reads sequences were demultiplexed using cutadapt (https://cutadapt.readthedocs.io/en/stable/) and aligned to the mm10 build of the mouse genome using snaptools (https://github.com/r3fang/SnapTools). Aligned bam files were converted to snap file format and then to sorted fragment files using snaptools’ snap-pre and dump-fragment functions respectively.

### Quality Control for snATAC-seq Data

Using ArchR (https://www.archrproject.com/index.html), a pipeline tool for the analysis of single-cell ATACseq data, arrow files were created for each sample only taking into account cells which had a minimum average transcription start site enrichment of 8 and a minimum of 1000 unique fragments. Doublet scores were added and doublets filtered based on default ArchR parameters. We created a gene score matrix based on a gene score model (using model 42 in this study^25^) that best correlated with gene-level chromatin accessibility to gene expression.

### Dimensionality Reduction and Initial Analyses of snATACseq data

Dimensionality reduction of the snATACseq data was carried out using ArchR’s addIterativeLSI functions with batch correction done using ArchR’s implementation of Harmony^60^, using default parameters in both functions. Following batch correction, nuclei from all samples were clustered using ArchR’s addClusters function with default parameters. The batch-corrected reduced dimensions were further projected for visualization on two dimensions via UMAP using ArchR’s addUMAP function with default parameters. Due to the large global differences between muscle fiber nuclei and mononuclear cell nuclei (compared to the differences among the mononuclear cells), it was difficult to further separate the different mononuclear cell types with fine resolution, on the same UMAP. Thus, we embedded the mononuclear cells on a new UMAP plane without the muscle fiber nuclei.

For downstream analysis, we focused on 54,000 nuclei from mononuclear cells. For these, we performed dimensionality reduction, sample and batch correction, and clustering using the same functions as above, with default parameters. We further projected the reduced dimension to two dimensions using ArchR’s addUMAP function, with a minimum distance of 0.3. Following UMAP projection, we annotated cell types based on the clusters which had maximum accessibility (based on calculated gene scores), for corresponding cell-type specific genes. Genome view of gene accessibility was done using ArchR’s ArchRBrowser function.

### Transcription Factor Enrichment Analysis

Transcription factors (TFs) from the Cisbp database (http://cisbp.ccbr.utoronto.ca/) were used for motif annotations. For motif enrichment, we used cell type-specific cREs and, using the one-sided Fisher’s exact test (implemented in ArchR), identified enriched TF motifs. An FDR cut-off of 0.1 and a log2FoldChange of 0.5 were used to identify significantly enriched TF motifs. Footprint and heatmap plots were generated using ArchR’s corresponding functions with default options, however the smoothWindow parameter for the plotFootprints function was set to 40.

### Trajectory analysis of snATAC-seq

Trajectory construction was done using ArchR’s AddTrajectory function, guided by previously identified clusters, biological time points and chromatin accessibility patterns for quiescence-associated and muscle differentiation-associated genes.

### Computing Cell Cycling Index

The gene list for calculating cell cycle index was obtained from a recent robust study by Giotti and colleagues^31^. We obtained an initial list of 701 identified cell-cycle regulating genes, sorted them based on three criteria: 1) their relevance in the G2M stage of the cell cycle, 2) genes previously known (before the Giotti et al. study) to play a role in cell cycle regulation, and 3) the number of previous studies citing the gene’s role in cell cycle regulation (selecting genes with previous citations of 9 and above). This resulted in a smaller set of 119 high-confidence G2M regulating genes which was used to calculate the cell cycle index using ArchR’s AddModuleScore function.

### Single-cell RNAseq data analysis

The fastq files for the external dataset from De Micheli et al was obtained from the sequence read archive (https://www.ncbi.nlm.nih.gov/sra, accession numbers: SRR10870296, SRR10870297, and SRR10870298). CellRanger (version 6.0.1) was used for generating count matrices from fastq files (both internally generated and external) and the filtered feature-barcode matrix was used to create a seurat object using Seurat (version 4.0.0). Counts were normalized using the SCTransform function while regressing out mitochondrial genes. Harmony was used to correct the source (external dataset versus internally generated) and sample effects, and subsequent downstream analyses were done using the Harmony-generated reduced dimension. The Pax7-expressing cluster was isolated and re-analyzed with logarithm normalization and similar batch correction approach using Harmony. Subsequent analysis (clustering etc) were done using the batch-corrected reduced dimension.

## SUPPLEMENTARY FIGURES AND LEGENDS

**Figure S1:**
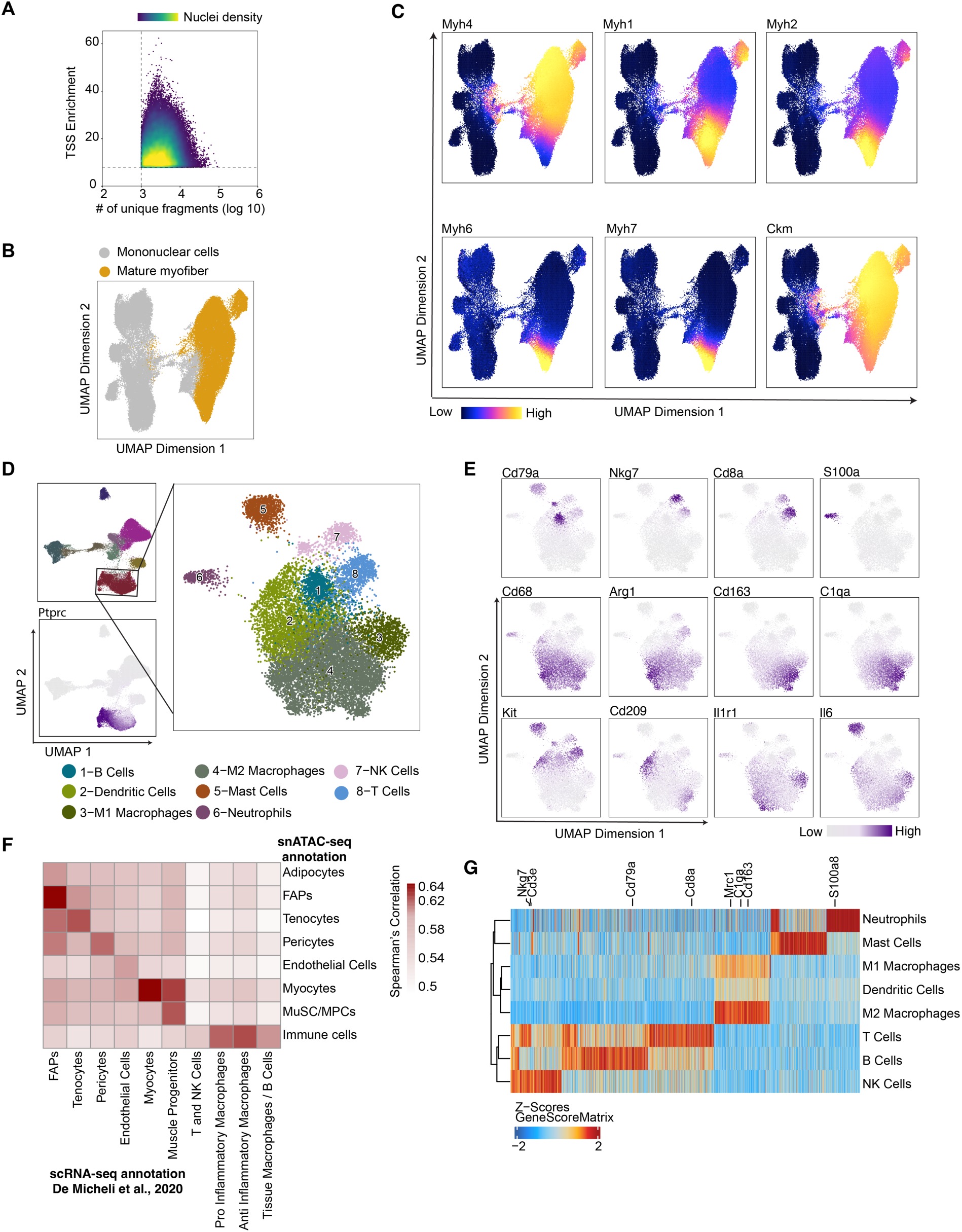
Single-nuclei ATAC-seq analysis of mouse skeletal muscle before and after multiple times of acute injury. **(A)** Scatter plot of the number of unique accessible fragments (x-axis, log10 transformed) and Tn5-insertion enrichment at transcription start sites (TSS, y-axis). A TSS enrichment threshold of 8 (horizontal dashed line) and a unique fragment number threshold of 1000 (vertical dashed line) were used for filtering nuclei resulting in 114,004 high-quality nuclei. **(B)** UMAP highlights the mature muscle fiber nuclei in yellow alongside nuclei from mononuclear cells of the muscle, including muscle progenitor cells. **(C)** On the UMAP including all 114,004 nuclei, the nuclei show very high gene scores of mature muscle fiber genes (the myosin heavy chain genes, *Acta1* and *Ckm*) are defined as mature myofiber nuclei. **(D)** UMAP plot distinguishes sub-populations of all the immune cells shown as cluster 4 in Fig 1B. **(E)** UMAP of sub-clustered immune cells are highlighted by the gene score of the marker genes of the various immune cell types identified in Fig S1D. **(F)** Heatmap of the global gene-to-gene Spearman’s correlation coefficient between major cell types identified from our in-house snATAC-seq data versus major cell types identified from an external single-cell RNA-seq dataset ^20^. **(G)** Heatmap of identified marker genes (based on gene scores) of immune cells (Wilcoxon Rank Sum test, false discovery rate FDR<=0.01 and Log2(Fold Change)>=0.5). Gene scores are scaled across cell types and cell types are clustered (hierarchical clustering).

**Figure S2:**
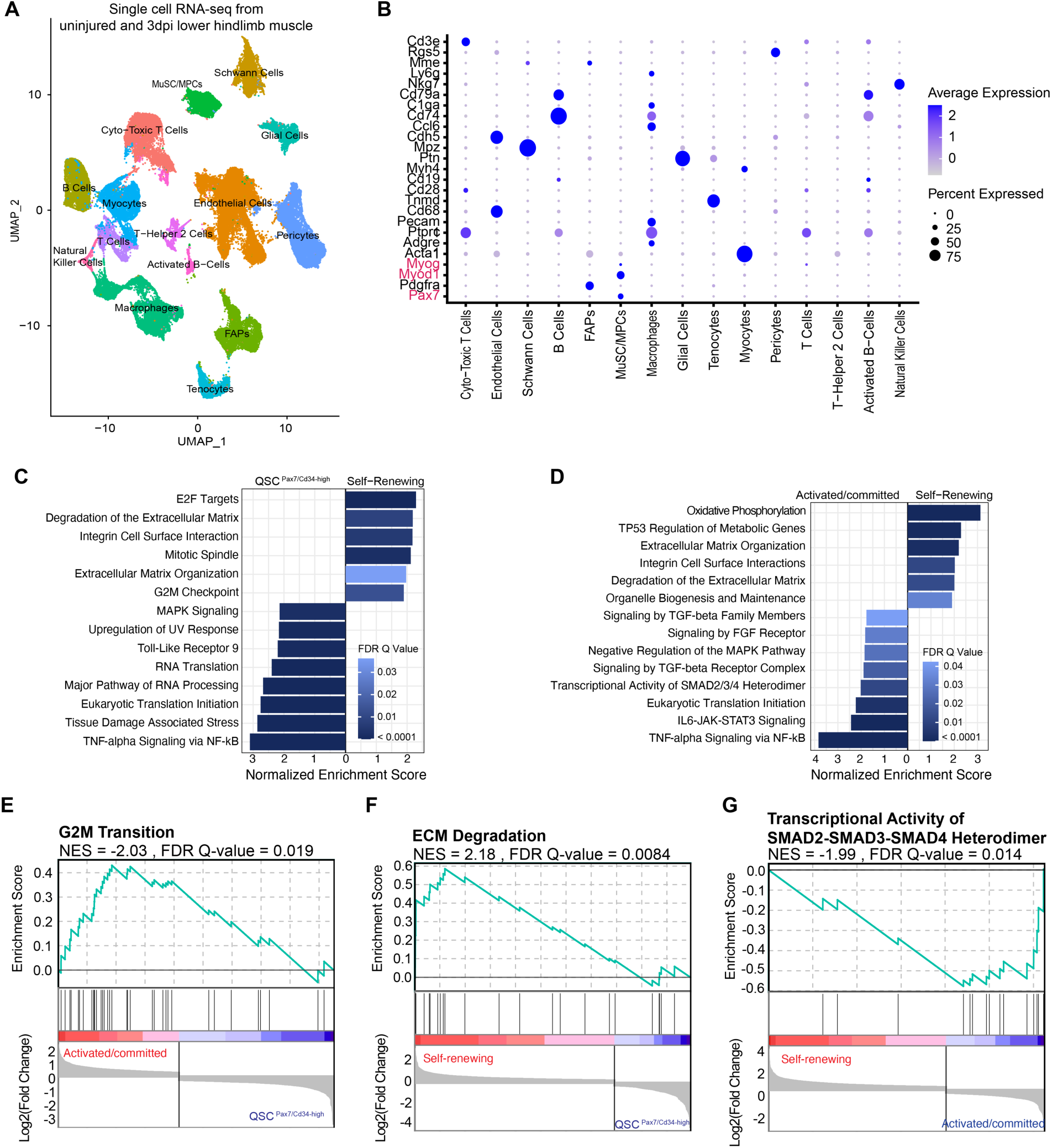
Single-cell RNA-seq analysis of the whole-cell isolates obtained from the lower hindlimb muscle of uninjured and 3dpi mice. **(A)** UMAP shows major cell types identified from single-cell RNA-seq analysis including both uninjured and injured (3 dpi) cells. **(B)** The expression pattern of the known marker genes of major cell types identified in Fig. S2A. The MuSC/MPC cells were used for sub-clustering analysis shown in Fig 3A. **(C, D)** Summary of Gene Set Enrichment Analysis (GSEA) results comparing QSC*^Pax7^*^/*Cd34*-high^ versus self-renewing MuSCs **(C)**, and Activated/Committed MPCs versus self-renewing MuSCs **(D)**. FDR is false discovery rate. **(E, F, G)** Specific GSEA plots. NES is normalized enrichment score.

**Figure S3:**
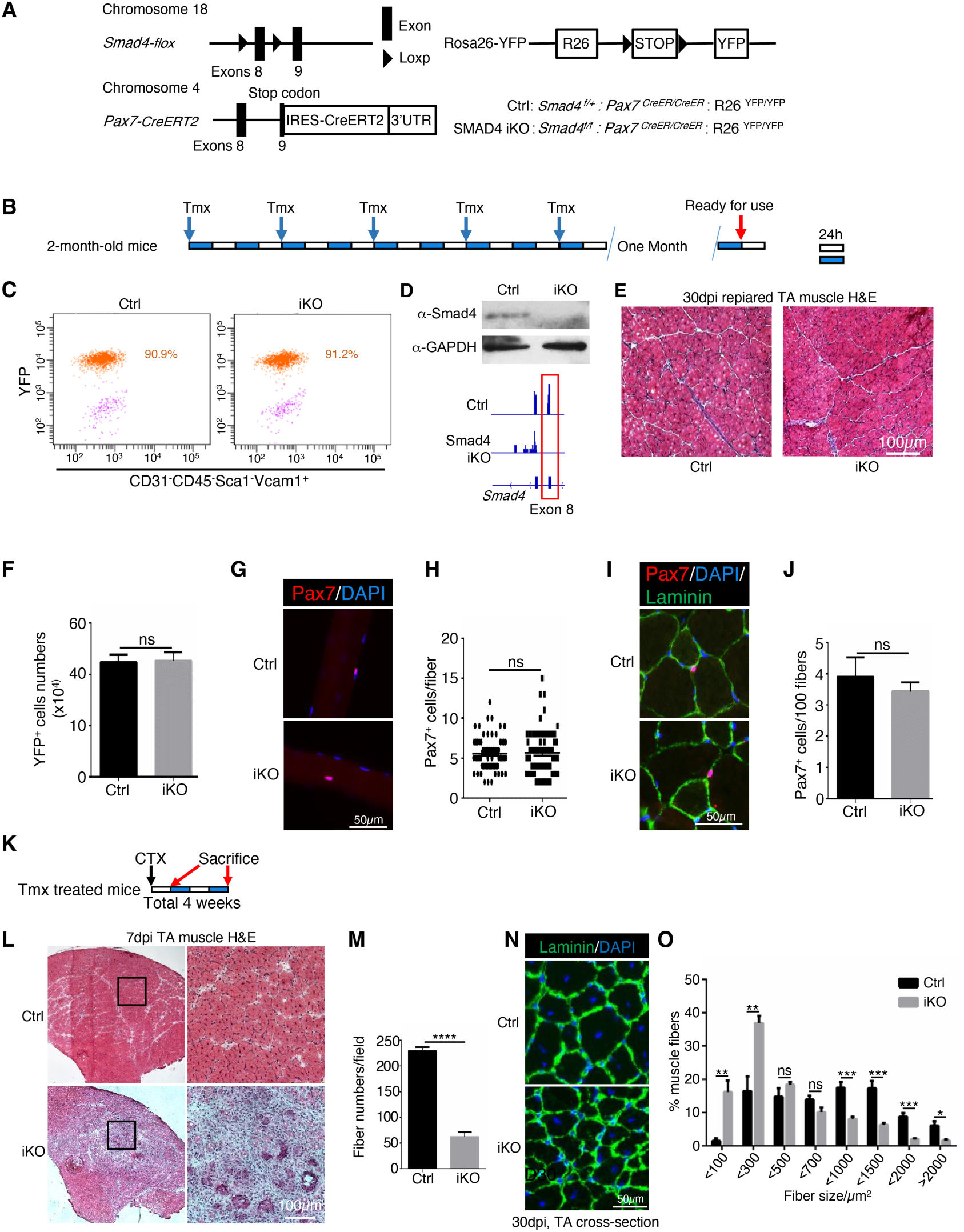
MuSC-specific SMAD4 KO does not affect adult MuSC maintenance but disrupts skeletal muscle regeneration. **(A)** Scheme of generation of *Smad4* MuSCs specific KO mice. **(B)** Scheme of Tamoxifen treatment of adult mice. Each white or blue bar represents 24 hours. **(C)** MuSCs from Tmx treated Ctrl and *Smad4* KO mice were sorted by FACS. The percentage of YFP^+^ cells in the MuSCs (CD31^-^CD45^-^Sca1^-^Vcam1^+^ cells) were counted, indicating the Cre recombination efficiency. **(D)** MuSCs sorted from Tmx treated Ctrl and KO mice were subjected to Westernblot analysis (top). (Bottom) 40-hour post CTX induced injury, the YFP+ MuSC/MPCs were FACS sorted from the injured lower hindlimb muscle of control and SMAD4 iKO mice. The collected cells were subjected to RNA-seq analysis. Representative RNA-seq profiles at the *Smad4* locus in MuSCs from 4 pairs of Ctrl and *Smad4* KO mice. The exons boxed in red at the *Smad4* locus were flanked by loxP and deleted in Tmx treated KO mice. **(E)** 30 days after tamoxifen (Tmx) administration, TA sections were subjected to H&E staining. The images are representative of multiple sections from 3 pairs of littermates. Scale bars, 100 µm. **(F)** MuSCs were sorted from Tmx-treated Ctrl and KO mice and the cell numbers were recorded. n=4. **(G)** Freshly isolated EDL fibers (n>30) from 3 pairs of mice were subjected to immunostaining for Pax7. The nuclei were counterstained with DAPI. The images are representative of multiple fibers. Scale bars, 50 µm. **(H)** Quantification of Pax7^+^ MuSCs number from (G). n=3. **(I)** Adult TA sections were subjected to immunostaining for Pax7 and Laminin. The nuclei were counterstained with DAPI. The images are representative of multiple sections from 3 pairs of littermates. Scale bars, 50 µm. **(J)** Quantification of Pax7^+^ cell number/100 myofibers from TA sections from 3 pairs of littermates in (I). **(K)** Schematic of the experimental design for muscle regeneration assay. Long range Tmx treated mice were injured by CTX at TA muscles. TAs were collected after 7 days or 30 days respectively and subjected to further experiments. **(L)** 7 days after CTX injury, TA sections were subjected to H&E staining. The images are representative of multiple sections from 6 pairs of mice. Scale bars, 100 µm. **(M)** Quantification of central nuclear fibers numbers from (L). n=6. The regenerated fiber numbers were counted on section images taken by a 20x microscope. **(N)** 30 days after CTX injury, TA sections were subjected to immunostaining for laminin. The nuclei were counterstained with DAPI. The images are representative of multiple sections from 3 pairs of littermates. Scale bars, 50 µm. **(O)** Quantification of the cross-section area (CSA) of individual myofibers from (N). The percentage of myofibers with a defined range of CSA over the total myofibers was calculated. Myofibers from 3 pairs of littermates were counted and more than 500 fibers from each pair of mice. All the results are presented as mean ± SEM. *p < 0.05, **p < 0.01, ***p < 0.001, ****p < 0.0001.

## SUPPLEMENTARY TABLES

**Table S1. Temporally-Resolved Cell Type-Specific Gene Score Across Muscle Regeneration Times** (Related to Figure 1D)

**Table S2. Temporally-Resolved Cell Type-Specific cRE Accessibility Across Muscle Regeneration Times** (Related to Figure 1E)

**Table S3. Temporally-Resolved Cell Type-Specific Motif Across Muscle Regeneration Times** (Related to Figure 1G)

**Table S4. Differentially Expressed Genes per MuSC/MPC sub-type in Muscle Regeneration** (Related to Figure 5)

**Table S5. Significant GSEA Terms of each sub-cluster of MuSC/MPCs** (Related to Figure 3)

**Table S6. Significantly Enriched Biological Terms Self-Renewing versus Genuine Quiescent MuSCs in Muscle Regeneration** (Related to Figure 5B)

**Table S7. DNA oligos used in this study for snATAC-seq and PCR.**

## REFERENCES

1. Brack, A. S. & Rando, T. A. Tissue-Specific Stem Cells: Lessons from the Skeletal Muscle Satellite Cell. Cell Stem Cell vol. 10 504–514 (2012).

2. Buckingham, M. & Relaix, F. The role of Pax genes in the development of tissues and organs: Pax3 and Pax7 regulate muscle progenitor cell functions. Annu. Rev. Cell Dev. Biol. 23, 645–673 (2007).

3. Murphy, M. & Kardon, G. Origin of vertebrate limb muscle: the role of progenitor and myoblast populations. Curr. Top. Dev. Biol. 96, 1–32 (2011).

4. Collins, C. A. et al. Stem Cell Function, Self-Renewal, and Behavioral Heterogeneity of Cells from the Adult Muscle Satellite Cell Niche. Cell 122, 289–301 (2005).

5. Yin, H., Price, F. & Rudnicki, M. A. Satellite cells and the muscle stem cell niche. Physiol. Rev. 93, 23–67 (2013).

6. Kassar-Duchossoy, L. et al. Pax3/Pax7 mark a novel population of primitive myogenic cells during development. Genes Dev. 19, 1426–1431 (2005).

7. Relaix, F., Rocancourt, D., Mansouri, A. & Buckingham, M. A Pax3/Pax7-dependent population of skeletal muscle progenitor cells. Nature vol. 435 948–953 (2005).

8. Tajbakhsh, S. Skeletal muscle stem cells in developmental versus regenerative myogenesis. J. Intern. Med. 266, 372–389 (2009).

9. Dumont, N. A., Bentzinger, C. F., Sincennes, M.-C. & Rudnicki, M. A. Satellite cells and skeletal muscle regeneration. Compr. Physiol. 5, 1027–1059 (2015).

10. Pawlikowski, B., Lee, L., Zuo, J. & Kramer, R. H. Analysis of human muscle stem cells reveals a differentiation-resistant progenitor cell population expressing Pax7 capable of self-renewal. Dev. Dyn. 238, 138–149 (2009).

11. Quarta, M. et al. An artificial niche preserves the quiescence of muscle stem cells and enhances their therapeutic efficacy. Nat. Biotechnol. 34, 752–759 (2016).

12. Feige, P. & Rudnicki, M. A. Isolation of satellite cells and transplantation into mice for lineage tracing in muscle. Nat. Protoc. 15, 1082–1097 (2020).

13. Beauchamp, J. R., Morgan, J. E., Pagel, C. N. & Partridge, T. A. Dynamics of Myoblast Transplantation Reveal a Discrete Minority of Precursors with Stem Cell–like Properties as the Myogenic Source. J. Cell Biol. 144, 1113–1122 (1999).

14. Qu, Z. et al. Development of approaches to improve cell survival in myoblast transfer therapy. J. Cell Biol. 142, 1257–1267 (1998).

15. Machado, L. et al. In Situ Fixation Redefines Quiescence and Early Activation of Skeletal Muscle Stem Cells. Cell Rep. 21, 1982–1993 (2017).

16. van Velthoven, C. T. J., de Morree, A., Egner, I. M., Brett, J. O. & Rando, T. A. Transcriptional Profiling of Quiescent Muscle Stem Cells In Vivo. Cell Rep. 21, 1994–2004 (2017).

17. van den Brink, S. C. et al. Single-cell sequencing reveals dissociation-induced gene expression in tissue subpopulations. Nat. Methods 14, 935–936 (2017).

18. Machado, L. et al. Tissue damage induces a conserved stress response that initiates quiescent muscle stem cell activation. Cell Stem Cell 28, 1125–1135.e7 (2021).

19. Oprescu, S. N., Yue, F., Qiu, J., Brito, L. F. & Kuang, S. Temporal Dynamics and Heterogeneity of Cell Populations during Skeletal Muscle Regeneration. iScience 23, 100993 (2020).

20. De Micheli, A. J. et al. Single-Cell Analysis of the Muscle Stem Cell Hierarchy Identifies Heterotypic Communication Signals Involved in Skeletal Muscle Regeneration. Cell Rep. 30, 3583–3595.e5 (2020).

21. Dell’Orso, S. et al. Single cell analysis of adult mouse skeletal muscle stem cells in homeostatic and regenerative conditions. Development 146, (2019).

22. Barruet, E. et al. Functionally heterogeneous human satellite cells identified by single cell RNA sequencing. Elife 9, e51576 (2020).

23. Preissl, S., Gaulton, K. J. & Ren, B. Characterizing cis-regulatory elements using single-cell epigenomics. Nat. Rev. Genet. (2022) doi:10.1038/s41576-022-00509-1.

24. Rodgers, J. T. et al. mTORC1 controls the adaptive transition of quiescent stem cells from G0 to G(Alert). Nature 510, 393–396 (2014).

25. Granja, J. M. et al. ArchR is a scalable software package for integrative single-cell chromatin accessibility analysis. Nat. Genet. 53, 403–411 (2021).

26. Gopinath, S. D., Webb, A. E., Brunet, A. & Rando, T. A. FOXO3 promotes quiescence in adult muscle stem cells during the process of self-renewal. Stem Cell Reports 2, 414–426 (2014).

27. Mourikis, P. et al. A critical requirement for notch signaling in maintenance of the quiescent skeletal muscle stem cell state. Stem Cells 30, 243–252 (2012).

28. Shea, K. L. et al. Sprouty1 regulates reversible quiescence of a self-renewing adult muscle stem cell pool during regeneration. Cell Stem Cell 6, 117–129 (2010).

29. Almada, A. E. & Wagers, A. J. Molecular circuitry of stem cell fate in skeletal muscle regeneration, ageing and disease. Nat. Rev. Mol. Cell Biol. 17, 267–279 (2016).

30. Bjornson, C. R. R. et al. Notch signaling is necessary to maintain quiescence in adult muscle stem cells. Stem Cells 30, 232–242 (2012).

31. Giotti, B. et al. Assembly of a parts list of the human mitotic cell cycle machinery. J. Mol. Cell Biol. 11, 703–718 (2019).

32. Chen, J.-F. et al. The role of microRNA-1 and microRNA-133 in skeletal muscle proliferation and differentiation. Nat. Genet. 38, 228–233 (2006).

33. Liu, L., Cheung, T. H., Charville, G. W. & Rando, T. A. Isolation of skeletal muscle stem cells by fluorescence-activated cell sorting. Nat. Protoc. 10, 1612–1624 (2015).

34. García-Prat, L. et al. FoxO maintains a genuine muscle stem-cell quiescent state until geriatric age. Nat. Cell Biol. 22, 1307–1318 (2020).

35. de Morrée, A. et al. Staufen1 inhibits MyoD translation to actively maintain muscle stem cell quiescence. Proc. Natl. Acad. Sci. U. S. A. 114, E8996–E9005 (2017).

36. Yue, L., Wan, R., Luan, S., Zeng, W. & Cheung, T. H. Dek Modulates Global Intron Retention during Muscle Stem Cells Quiescence Exit. Dev. Cell 53, 661–676.e6 (2020).

37. Motohashi, N. & Asakura, A. Muscle satellite cell heterogeneity and self-renewal. Front Cell Dev Biol 2, 1 (2014).

38. Sincennes, M.-C., Brun, C. E. & Rudnicki, M. A. Concise Review: Epigenetic Regulation of Myogenesis in Health and Disease. Stem Cells Transl. Med. 5, 282–290 (2016).

39. Vander Ark, A., Cao, J. & Li, X. TGF-β receptors: In and beyond TGF-β signaling. Cell. Signal. 52, 112–120 (2018).

40. Lowery, J. & Caestecker, M. de. Bmp Signaling and Vascular Disease. in Encyclopedia of Biological Chemistry (Second Edition) (eds. Lennarz, W. J. & Lane, M. D.) 229–239 (Academic Press, 2013).

41. Pawlak, J. B. & Blobe, G. C. TGF-β superfamily co-receptors in cancer. Dev. Dyn. 251, 137–163 (2022).

42. Budi, E. H., Duan, D. & Derynck, R. Transforming Growth Factor-β Receptors and Smads: Regulatory Complexity and Functional Versatility. Trends Cell Biol. 27, 658–672 (2017).

43. Robinson, D. C. L. et al. Negative elongation factor regulates muscle progenitor expansion for efficient myofiber repair and stem cell pool repopulation. Dev. Cell 56, 1014–1029.e7 (2021).

44. Bi, P. et al. Control of muscle formation by the fusogenic micropeptide myomixer. Science vol. 356 323–327 (2017).

45. Millay, D. P. et al. Myomaker is a membrane activator of myoblast fusion and muscle formation. Nature 499, 301–305 (2013).

46. Lingbeck, J. M., Trausch-Azar, J. S., Ciechanover, A. & Schwartz, A. L. In vivo interactions of MyoD, Id1, and E2A proteins determined by acceptor photobleaching fluorescence resonance energy transfer. FASEB J. 22, 1694–1701 (2008).

47. Partridge, T. A., Grounds, M. & Sloper, J. C. Evidence of fusion between host and donor myoblasts in skeletal muscle grafts. Nature 273, 306–308 (1978).

48. Massagué, J. A very private TGF-beta receptor embrace. Molecular cell vol. 29 149–150 (2008).

49. Yanagisawa, M., Mukai, A., Shiomi, K., Song, S.-Y. & Hashimoto, N. Community effect triggers terminal differentiation of myogenic cells derived from muscle satellite cells by quenching Smad signaling. Exp. Cell Res. 317, 221–233 (2011).

50. Ono, Y. et al. BMP signalling permits population expansion by preventing premature myogenic differentiation in muscle satellite cells. Cell Death Differ. 18, 222–234 (2010).

51. Cohen, T. V., Kollias, H. D., Liu, N., Ward, C. W. & Wagner, K. R. Genetic disruption of Smad7 impairs skeletal muscle growth and regeneration. J. Physiol. 593, 2479–2497 (2015).

52. Lamarche, É. et al. SMAD2 Promotes Myogenin Expression and Terminal Myogenic Differentiation. bioRxiv 2020.07.28.225888 (2020) doi:10.1101/2020.07.28.225888.

53. Paris, N. D., Soroka, A., Klose, A., Liu, W. & Chakkalakal, J. V. Smad4 restricts differentiation to promote expansion of satellite cell derived progenitors during skeletal muscle regeneration. Elife 5, (2016).

54. Stantzou, A. et al. BMP signaling regulates satellite cell-dependent postnatal muscle growth. Development 144, 2737–2747 (2017).

55. Liu, D., Black, B. L. & Derynck, R. TGF-beta inhibits muscle differentiation through functional repression of myogenic transcription factors by Smad3. Genes Dev. 15, 2950–2966 (2001).

56. Liu, D., Kang, J. S. & Derynck, R. TGF-beta-activated Smad3 represses MEF2-dependent transcription in myogenic differentiation. EMBO J. 23, 1557–1566 (2004).

57. Dong, Y., Lakhia, R. & Thomas, S. S. Interactions between p-Akt and Smad3 in injured muscles initiate myogenesis or fibrogenesis. American Journal (2013).

58. Ge, X. et al. Smad3 signaling is required for satellite cell function and myogenic differentiation of myoblasts. Cell Res. 21, 1591–1604 (2011).

59. Ge, X. et al. Lack of Smad3 signaling leads to impaired skeletal muscle regeneration. Am. J. Physiol. Endocrinol. Metab. 303, E90–102 (2012).

60. Korsunsky, I. et al. Fast, sensitive and accurate integration of single-cell data with Harmony. Nat. Methods 16, 1289–1296 (2019).

